# Neto-α controls synapse organization and homeostasis at the *Drosophila* neuromuscular junction

**DOI:** 10.1101/812040

**Authors:** Tae Hee Han, Rosario Vicidomini, Cathy Isaura Ramos, Qi Wang, Peter Nguyen, Michal Jarnik, Moyi Li, Michal Stawarski, Roberto X. Hernandez, Gregory T. Macleod, Mihaela Serpe

## Abstract

Glutamate receptor auxiliary proteins control receptor distribution and function, ultimately controlling synapse assembly, maturation and plasticity. At the *Drosophila* neuromuscular junction (NMJ), a synapse with both pre- and post-synaptic kainate-type glutamate receptors (KARs), we show that the auxiliary protein Neto evolved functionally distinct isoforms to modulate synapse development and homeostasis. Using genetics, cell biology and electrophysiology we demonstrate that Neto-α functions on both sides of the NMJ. In muscle, Neto-α limits the size of the postsynaptic receptors field. In motor neurons, Neto-α controls neurotransmitter release in a KAR-dependent manner. Furthermore, Neto-α is both required and sufficient for the presynaptic increase in neurotransmitter release in response to reduced postsynaptic sensitivity. This KAR-independent function of Neto-α is involved in activity-induced cytomatrix remodeling. We propose that *Drosophila* ensured NMJ functionality by acquiring two Neto isoforms with differential expression patterns and activities.

## Introduction

Formation of functional synapses during development, and their fine-tuning during plasticity and homeostasis relies on ion channels and their accessory proteins, which control where, when and how the channels function. Auxiliary proteins are diverse transmembrane proteins which associate with channel complexes at all stages of their life-cycle and mediate their properties as well as subcellular distribution, surface expression, synaptic recruitment, and associations with various synaptic scaffolds (Jackson and Nicoll, 2011). Channel subunits expanded and diversified during evolution to impart different channel biophysical properties (Alberstein et al., 2015; Han et al., 2015; Li et al., 2016; Mayer, 2017), but whether auxiliary proteins evolved to match channel diversity remains unclear.

Ionotropic glutamate receptors (iGluRs) mediate neurotransmission at most excitatory synapses in the vertebrate CNS and at the neuromuscular junction (NMJ) of insects and crustaceans and include α-amino-3-hydroxy-5-methyl-4-isoxazolepropionic acid (AMPA), N-methyl-D-aspartic acid (NMDA), and kainate (KA) receptors. Sequence analysis of the *Drosophila* genome identified 14 iGluRs genes that resemble vertebrate AMPA, NMDA and KA receptors (Littleton and Ganetzky, 2000). The fly receptors have strikingly different ligand binding profiles (Han et al., 2015; Li et al., 2016); nonetheless, phylogenetic analysis indicate that two of the *Drosophila* genes code for AMPA receptors, two for NMDAR, and 10 for subunits of the KAR family, which is highly expanded in insects (Li et al., 2016). In flies as in vertebrates, AMPA and KA receptors have conserved, dedicated auxiliary proteins. For example, AMPARs rely on Stargazin and its relatives which selectively modulate receptors gating properties, trafficking, and interactions with scaffolds such as PSD-95-like membrane-associated guanylate kinases (Milstein and Nicoll, 2008; Sumioka et al., 2010; Tomita et al., 2005; Tomita et al., 2003; Twomey et al., 2016); Stargazin is also required for the functional reconstitution of invertebrate AMPARs (Li et al., 2016; Walker et al., 2006). KARs are modulated by Neto (<u>Ne</u>uropilin and <u>To</u>lloid-like) family of proteins, that include the vertebrate Neto1 and −2 (Ng et al., 2009; Zhang et al., 2009), *C. elegans* SOL-2/Neto (Wang et al., 2012) and *Drosophila* Neto (Kim et al., 2012; Kim and Serpe, 2013). Neto proteins differentially modulate the gating properties of vertebrate KARs (Tomita and Castillo, 2012). A role for Neto in the biology of KARs *in vivo* has been more difficult to assess because of the low levels of KARs and Neto proteins (Lerma and Marques, 2013). Nevertheless, vertebrate Netos could modulate the synaptic recruitment of selective KARs by association with synaptic scaffolds such as GRIP and PSD-95; in fact, the PDZ-binding domains of vertebrate KAR/Neto complexes are essential for basal synaptic transmission and LTP (Sheng et al., 2018; Tang et al., 2012). Post-translational modifications regulate Neto activities *in vitro*, but the *in vivo* relevance of many of these observations remains unknown (Lomash et al., 2017).

*Drosophila* NMJ is an excellent genetic system to probe the repertoire of Neto functions. This glutamatergic synapse appears to rely exclusively on KARs, with five postsynaptic subunits and one presynaptic subunit (see below). We have previously found that *Drosophila* Neto is an obligatory auxiliary subunit of the postsynaptic KAR complexes (Kim et al., 2012; Kim and Serpe, 2013): In the absence of Neto the postsynaptic KARs fail to cluster at synaptic sites and the animals die as completely paralyzed embryos. Heterologous reconstitution of postsynaptic KARs in *Xenopus* oocytes revealed that Neto is absolutely required for functional receptors (Han et al., 2015). The fly NMJ contains two receptor complexes (type-A and -B) with different subunits composition (either GluRIIA or -IIB, plus -IIC, -IID and -IIE), and distinct properties, regulation, and localization patterns (DiAntonio, 2006; DiAntonio et al., 1999; Featherstone et al., 2005; Marrus et al., 2004; Petersen et al., 1997; Qin et al., 2005). The postsynaptic response to the fusion of single synaptic vesicles (quantal size) is much reduced for NMJs with type-B receptors only; in fact, the dose of synaptic GluRIIA and GluRIIB is a key determinant of quantal size (DiAntonio et al., 1999). The fly NMJ is also a powerful model system to study homeostatic plasticity (Davis and Muller, 2015; Frank, 2014): Manipulations that decrease the responsiveness of postsynaptic glutamate receptor (leading to a decrease in quantal size) trigger a robust compensatory increase in presynaptic neurotransmitter release (quantal content) (Davis et al., 1998; DiAntonio et al., 1999; Petersen et al., 1997). This increase in quantal content restores evoked muscle responses to normal levels. A presynaptic KAR, KaiRID, has been recently implicated in basal neurotransmission and presynaptic homeostatic potentiation (PHP) at the larval NMJ (Kiragasi et al., 2017; Li et al., 2016). The role of KaiRID in modulation of basal neurotransmission resembles GluK2/GluK3 function as autoreceptors (Pinheiro et al., 2007). The role of KaiRID in PHP must be indirect since a mutation that renders this receptor Ca^2+^-impermeable has no effect on the expression of presynaptic homeostasis (Kiragasi et al., 2017). The fly NMJ reliance on KARs raises the possibility that *Drosophila* diversified and maximized its use of Neto proteins. *Drosophila neto* codes for two isoforms (Neto-α and Neto-β) with distinct intracellular domains generated by alternative splicing (Ramos et al., 2015); both cytoplasmic domains are rich in phosphorylation sites and docking motifs, suggesting rich modulation of Neto/KARs distribution and function. Indeed, Neto-β, the predominant isoform at the larval NMJ, mediates intracellular interactions that recruit PSD components and enables synaptic stabilization of selective receptor subtypes (Ramos et al., 2015). Neto-α can rescue viability and receptor clustering defects of *neto^null^* (Kim et al., 2012; Kim et al., 2015; Ramos et al., 2015). However, the endogenous functions of Neto-α remain unknown.

Here, we showed that Neto-α is key to synapse development and homeostasis and fulfils functions that are completely distinct from those of Neto-β. Using isoform specific mutants and tissue specific manipulations, we found that loss of Neto-α in the postsynaptic muscle disrupts glutamate receptor fields and produces enlarged PSDs. Loss of presynaptic Neto-α disrupts basal neurotransmission and renders these NMJs unable to express PHP. We mapped the different functions of Neto-α to distinct protein domains and demonstrated that Neto-α is both required and sufficient for PHP, functioning as a bona fide effector for PHP. We propose that *Drosophila* ensured NMJ functionality by acquiring two Neto isoforms with differential expression patterns and activities.

## Results

### Neto-α and Neto-β have distinct roles during NMJ development

To study the function of Neto-α at the *Drosophila* NMJ, we generated isoform specific *neto-α^null^* mutants using the CRISPR/Cas-9 technology (Figure 1A and Material and Methods). Several independent lines were isolated and confirmed molecularly as *neto-α* genetic null mutants; all these lines were viable, fertile and exhibited no obvious behavior deficits. For further analyses, we selected a line in which a total of 13,476 bp have been deleted, including the α-specific exon and parts of the flanking introns. To test whether this deletion affected the expression of Neto-β we measured the levels of *neto*-β transcript by qPCR in larval carcasses (not shown) and the levels of net Neto-β protein in the larval muscle by Western blot (Figure 1B) (Ramos et al., 2015). No changes were observed indicating that muscle expression of Neto-β was not affected by eliminating the α-specific exon. We tested whether Neto-β is properly targeted at *neto-α^null^* NMJs using anti-Neto antibodies raised against the extracellular CUB1 domain, common to both Neto isoforms (Kim et al., 2012) (Figures 1C and 1D). Quantification of these NMJ signals relative to anti-HRP, which labels neuronal membranes (Jan and Jan, 1982), confirmed our previous findings that Neto-β is the predominant isoform at the fly NMJ (Ramos et al., 2015) and indicated relatively normal synaptic recruitment of Neto-β in the absence of Neto-α.

**Figure 1:**
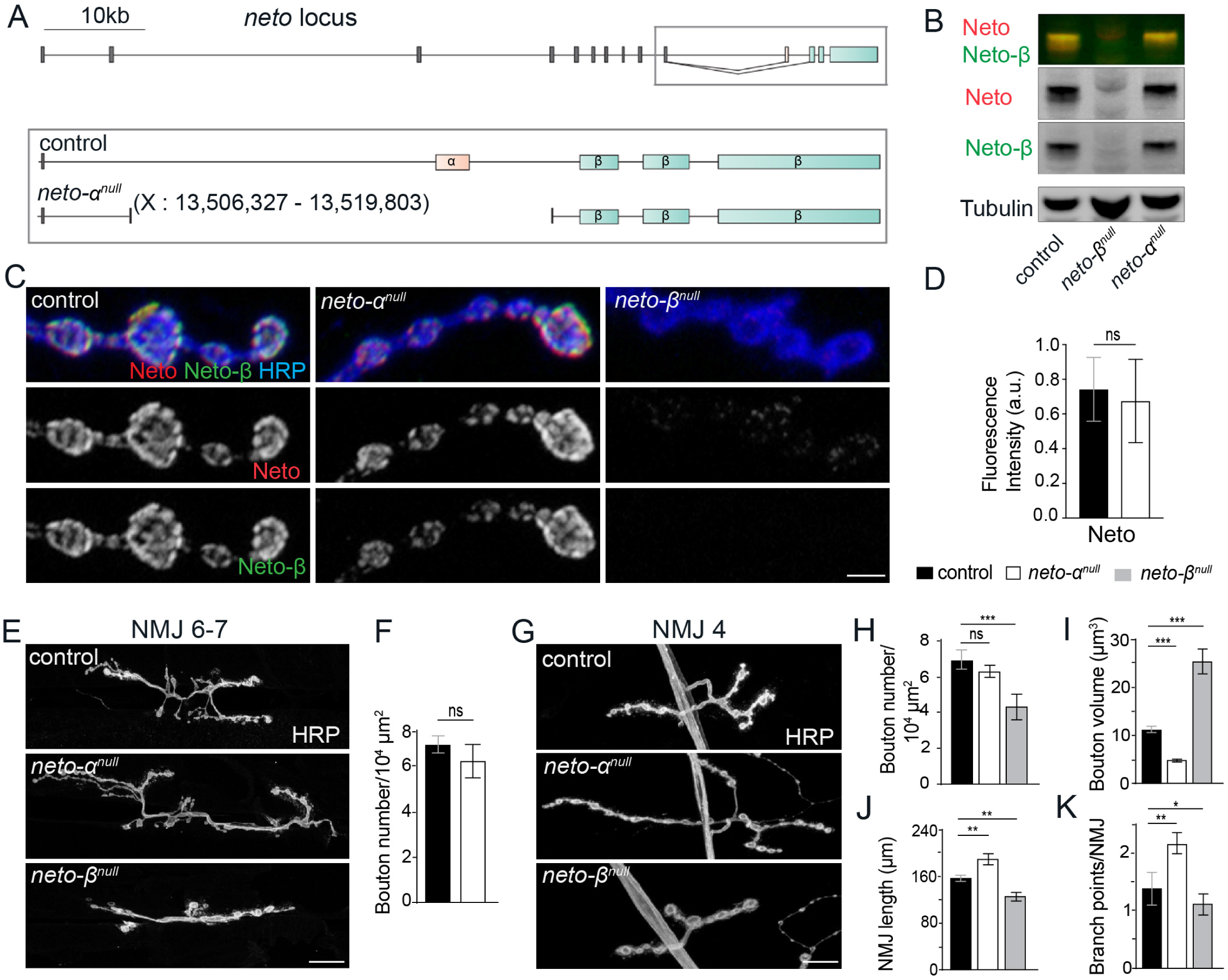
Neto-α has different functions than Neto-β at the NMJ. (A) Diagram of the *Drosophila neto* locus consisting of 10 shared exons coding for the extracellular and transmembrane domains (dark gray), an exon encoding Neto-α intracellular part (pink) and 3 exons coding for Neto-β cytoplasmic part (blue). (B) Western blot analysis of muscle extracts from control (*w^1118^*), *neto-α^null^* and *neto-β^null^* larvae labeled with anti-Neto (green), -Neto-β (red) and -Tubulin antibodies. Similar Neto-β levels were detected in control and *neto-α^null^*. (C) Confocal images of synaptic boutons (NMJ4/ segment A3) of indicated genotypes stained for Neto (red), Neto-β (green) and HRP (blue). *neto-α^null^* boutons show normal levels of Neto as quantified in (D); *neto-β^null^* boutons are shown for comparison. (E-K) Confocal images (E) and (G) and morphometric quantifications (F) and (H-K) of NMJ6-7 and NMJ4 (segment A3) in larvae of indicated genotypes. *neto-α^null^* NMJs have normal number (F and H) but smaller (I) boutons, and increased NMJ length (J) and branch points (K). Scale bars: 3µm (C) and 20µm (E, G). Data are represented as mean ± SEM.***, *p*<0.001; **, *p*<0.005; ns, *p*>0.05.

We have previously reported that loss of Neto-β alters the NMJ morphology and produces shorter NMJs with fewer, enlarged type-Ib boutons (Ramos et al., 2015). The morphology of *neto-α^null^* NMJs was strikingly different, with significantly smaller type-Ib boutons (NMJ6/7 and NMJ4 analyzed in Figures 1E-K). In the absence of Neto-α, the length of individual NMJ segments didn’t change significantly, but the number of branches increased, producing longer NMJs. Thus, the two Neto isoforms appear to have distinct roles during NMJ growth and development.

### Neto-α is required for normal NMJ physiology

To test whether Neto-α influences NMJ function, we recorded spontaneous miniature potentials (mEJPs) and evoked excitatory junctional potentials (EJPs) from muscle 6, segment A3, of third instar larvae of control (*w^1118^*) and *neto-α^null^* animals (Figures 2A-E, and Supplemental Table 1). No differences were found in the resting potential and input resistance in mutant larvae. The mini amplitude or quantal size, which reflects the amount of glutamate released from a single vesicle and the status of postsynaptic receptors, was slightly reduced in *neto-α^null^* animals compared to the control (*neto-α^null^*, 1.25 ± 0.05 mV vs. control - *w^1118^*, 1.09 ± 0.06 mV, *p* = 0.07, Figures 2A-B). This is different from *neto*-*β^null^* animals, which have significantly reduced postsynaptic type-A receptors and decreased quantal size (Ramos et al. 2015). However, *neto*-*β^null^* mutants have normal EJP amplitude, whereas *neto-α^null^* animals showed EJP amplitude reduced by half (19.75 ± 1.63, compared to 31.31 ± 2.40 in control, *p <* 0.0001, Figures 2C-D). The quantal content, estimated as ratio of average EJP amplitude to the mEJP amplitude, was decreased in *neto-α^null^* larvae (19.75 ± 1.63, compared to 31.31 ± 2.40 in control, *p*= 0.0008, Figure 2E). In contrast, the *neto*-*β^null^* mutants exhibit a robust compensatory increase in quantal content (Ramos et al. 2015), highlighting the differences between the two Neto isoforms at the *Drosophila* NMJ.

**Figure 2:**
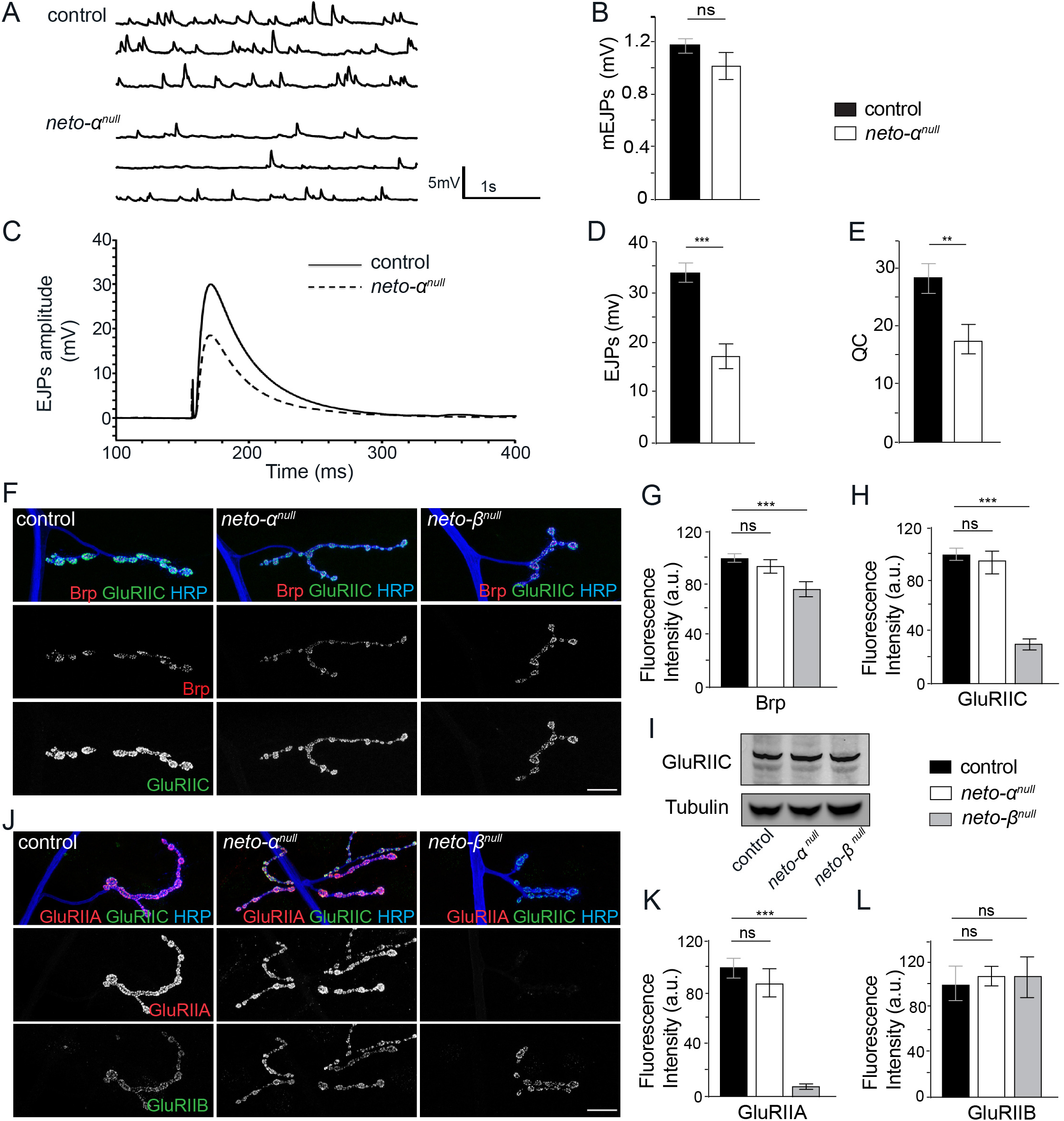
Neto-α is required for normal neurotransmitter release. (A-E) Representative traces of spontaneous (A) and evoked (C) neurotransmitter release recorded from muscle 6 of control (*w^1118^*) and *neto-α^null^* third instar larvae at 0.5 mM Ca^2+^. Summary bar graphs showing the mean amplitude of mEJPs (B), the mean amplitude EJPs (D), and the quantal content (QC)(E). *neto-α^null^* animals have normal mEJPs amplitude (B) but show reduced EJPs amplitude and QC (C). The numbers of NMJs recorded and the muscle resistance are indicated in Supplemental Table 1. (F) Confocal images of NMJ4 in larvae of indicated genotypes stained for Brp (red), GluRIIC (green) and HRP (blue). As quantified in (G-H), Brp and GluRIIC signals are normal in *neto-α^null^* animals, albeit severely reduced in *neto-β^null^* mutants, used here for comparison. (I) Western blot analysis shows normal GluRIIC muscle expression in both isoform specific alleles. (J) Confocal images of NMJ4 labeled for GluRIIA (red), GluRIIB (green) and HRP (blue). *neto-α^null^* animals have GluRIIA and GluRIIB levels similar to control, as quantified in (K-L). Scale bars: 20µm. Data are represented as mean ± SEM.***, *p*<0.001; **, *p*<0.005; ns, *p*>0.05.

### *neto-α^null^* animals have normal receptor levels but enlarged PSDs

Since Neto is key to the synaptic recruitment of postsynaptic KARs, we asked whether the defects observed at *neto-α^null^* NMJs are due to altered distribution of synaptic receptors. We first examined the synaptic distribution of GluRIIC, an essential subunit shared by both type-A and -B receptors, and of the presynaptic scaffold Bruchpilot (Brp), the fly homolog of the vertebrate active zone protein ELSK/CAST, which marks the sites of neurotransmitter release (Kittel et al., 2006; Marrus et al., 2004). The GluRIIC and Brp synaptic signals were in perfect juxtaposition at *neto-α^null^* NMJs (Figure 2F); the puncta appeared less intense in the absence of Neto-α (see below), but the relative levels of synaptic GluRIIC, as well as net GluRIIC protein in the larval muscle were normal (Figures 2G-I). This is in contrast to *neto*-*β^null^* or *neto^hypo^* mutants which have normal net levels of receptors in the larval muscle but severely reduced synaptic receptors, presumably because of limiting Neto (Kim et al., 2012; Ramos et al., 2015). Furthermore, in the absence of Neto-α, we could not detect any perturbations in the levels of synaptic GluRIIA or GluRIIB, demonstrating that Neto-α does not influence their synaptic recruitment (Figures 2J-L). This result is consistent with the normal mEJPs amplitude observed at *neto-α^null^* NMJs.

The mildly reduced GluRIIC signal intensities may indicate alterations in the size and/or organization of receptor fields. We tested this possibility by examining individual PSDs. In *Drosophila*, the PSD-95 ortholog Discs Large (Dlg) does not colocalize with the iGluR fields and instead is adjacent to the PSDs (Guan et al., 1996). Indeed, the boundaries between GluRIIC and Dlg marked structures were very well defined in control boutons but were no longer recognizable at *neto-α^null^* NMJs (Figure 3A-B). Moreover, the 3D-reconstructions of these boutons showed no overlap between GluRIIC and Dlg signals in controls, but significant overlap in *neto-α^null^* mutants.

To further characterize this defect, we examined synapses stained for pre- and postsynaptic components using 3D structured illumination microscopy (3D-SIM). The individual synapses were stained with Brp, which accumulates at presynaptic specializations called T-bars (Wagh et al., 2006). The anti-Brp monoclonal antibody NC82 recognizes an epitope on the outer diameter of the T-bars and produces a ring-shaped signal when examined by super-resolution microscopy (Fouquet et al., 2009; Sulkowski et al., 2016). Opposite to the T-bars, the PSDs contain the iGluR/Neto complexes stabilized by various postsynaptic proteins (Sulkowski et al., 2016). At *neto-α^null^* synapses the Brp rings appeared normal (Figures 3C-D, top panels), but the GluRIIC and Neto signals spread outward, expanding the boundaries of individual PSDs. 3D-reconstructions captured the enlarged receptor fields, which appeared to fill the small *neto-α^null^* boutons (Figures 3C-D, lower panels). To quantify these differences in PSD organization, we examined the individual synapses in serial section electron micrographs (see Materials and Methods, Figures 3E-H). The maximum diameters observed at mutant PSDs were significantly higher than the controls (1,100 nm in *neto-α^null^* vs. 780 nm in control, *w^1114^*). In contrast, the *neto-α^null^* T-bars appeared similar to those of control synapses. These results are consistent with our immunohistochemistry results and indicate that Neto-α limits the size of the postsynaptic receptor fields but has no detectable role in the organization of presynaptic specializations.

**Figure 3:**
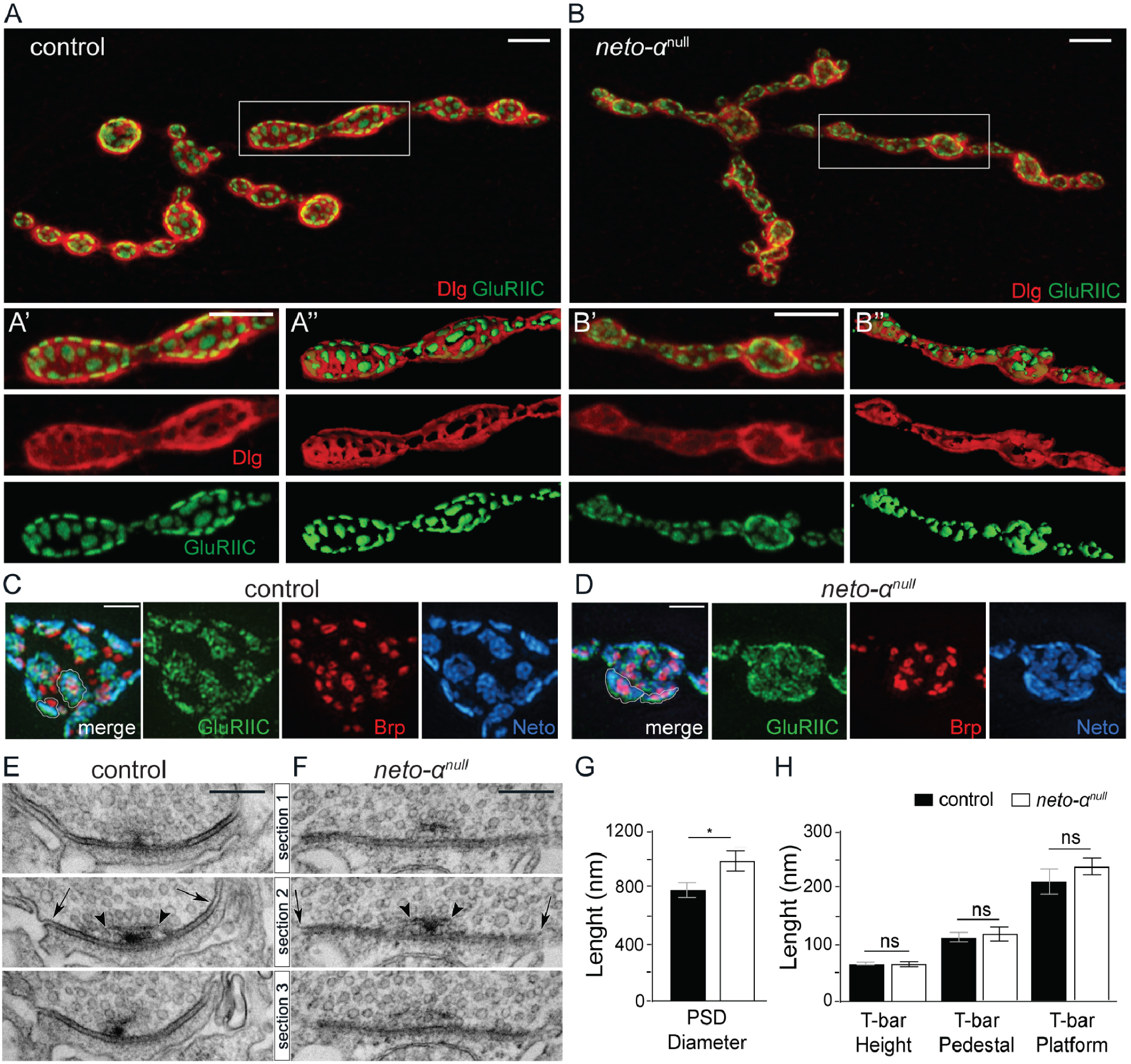
Neto-α limits the postsynaptic receptor fields. (A) Confocal images of NMJ4 labeled with Dlg (red) and GluRIIC (green). In control animals, Dlg-positive staining abuts on GluRIIC-marked PSDs (detail A’ was used for 3D-reconstruction in A’’). The borders between Dlg and GluRIIC are blurred in *neto-α^null^* boutons (B-B’’). (C-D) 3D-SIM images of NMJ4 boutons labeled with Brp (red), GluRIIC (green) and Neto (blue). Individual PSDs are clearly separated in control boutons but are difficult to distinguish in *neto-_αnull_*. (E-F) Serial sections of electron micrographs of single PSDs in control (E) and *neto-α^null^* boutons (F). The longest diameters detectable in serial sections for each PSD or T-bar structure are indicated by arrows and arrowheads, respectively (E-F), and are quantified (G-H). The numbers of PSDs examined: 10 control and 14 *neto-α^null^*; and T-bars: 8 control and 12 *neto-α^null^*. Scale bars: 10µm (A-B), 1µm (C-D). Data are represented as mean ± SEM.*, *p*<0.01; ns, p>0.05.

### Neto-α functions in both pre- and post-synaptic compartments

We next asked whether Neto-α activities are restricted to the postsynaptic compartment using tissue specific rescue and knockdown experiments. In these assays, the PSD areas were quantified from stacks of 3D-SIM images (as described in Materials and Methods). We found that expression of *neto-α* in motor neurons did not rescue the PSD sizes of *neto-α^null^* synapses, which remained enlarged, as seen in mutant PSDs (0.657 ± 0.004 µm^2^ in neuronal rescue (n=1569) vs 0.654 ± 0.004 µm^2^ in *neto-α^null^* (n=1438) (Figures 4A-C, quantified in 4E). However, muscle overexpression of a *neto-α* transgene fully rescued the PSD size of *neto-α^null^* synapses to a mean indistinguishable from control (0.570 ± 0.003 µm^2^ in muscle rescue (n=1600) vs. 0.580 ± 0.004 µm^2^ in control *(w^1114^*) (n=1487)) (Figures 4A, 4D and 4E). Even though the PSD sizes were variable, their relative frequency distribution showed that *neto-α^null^* PSDs were consistently larger than the control (Figure 4F); this was also captured by the right shifted cumulative frequency distribution of the observed *neto-α^null^* PSDs (Figure 4G). Once again, the distribution of neuron rescue PSDs was similar to that of *neto-α^null^* mutants, whereas the muscle rescue PSDs resembled the distribution of control PSDs. This indicates that Neto-α functions in the muscle to limit postsynaptic receptor fields. This conclusion was also supported by knockdown experiments (not shown).

**Figure 4:**
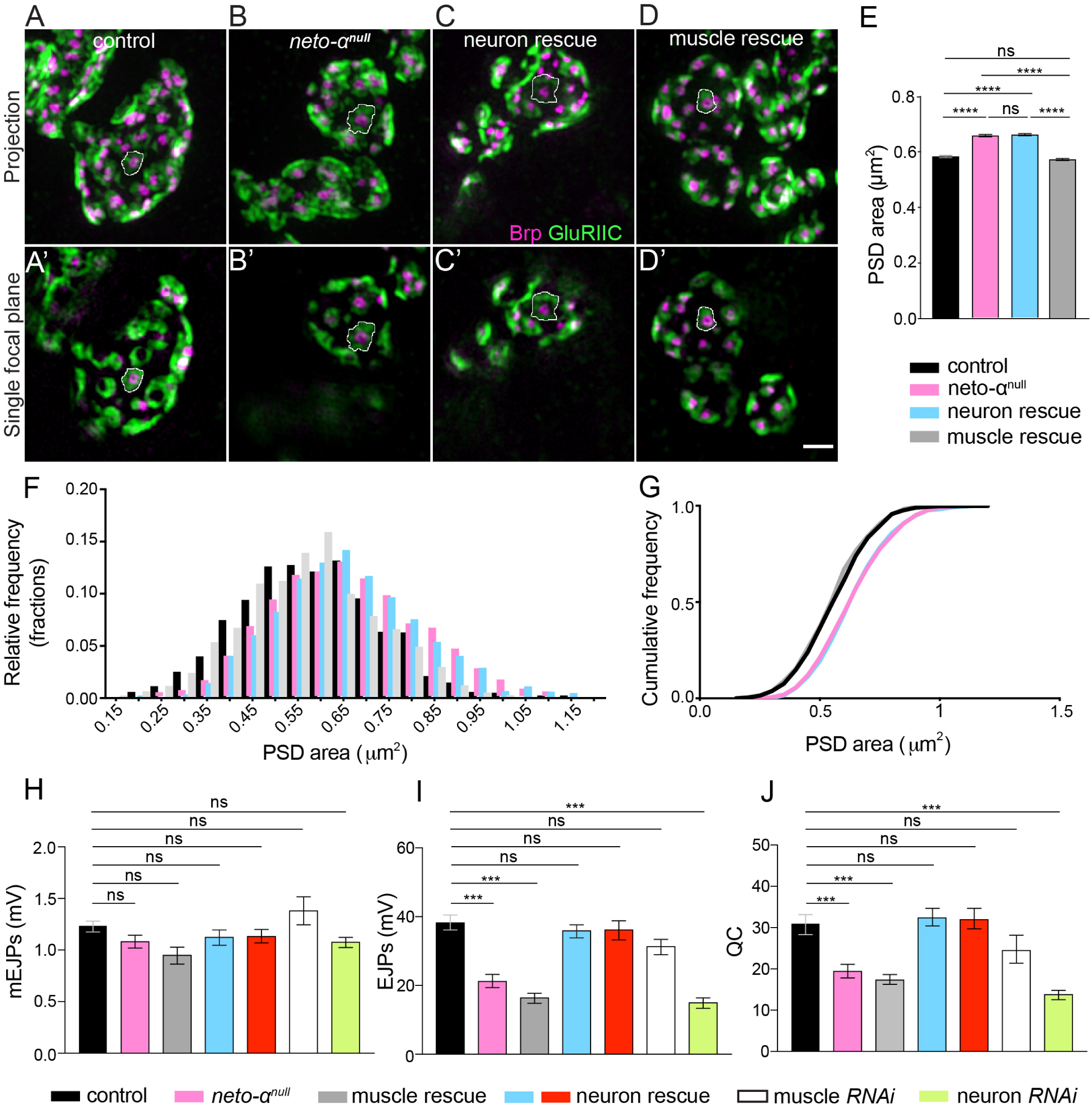
Neto-α functions in both pre- and post-synaptic compartments. (A-D) Representative 3D-SIM images (maximum intensity projection and single focal plane) of NMJ4 boutons of indicated genotypes labeled with Brp (magenta) and GluRIIC (green). Mean individual PSD areas (white contours) are plotted in (E). Muscle but not neuronal expression of Neto-α rescues the enlarged PSDs size of *neto-α^null^*. (F-G) Relative and cumulative frequency distribution of different size PSDs. 1400 or more individual PSDs from 12 NMJs were quantified for each genotype. (H-J) Summary bar graphs showing the mean amplitude of mEJPs (H), the mean amplitude EJPs (I), and the quantal content (QC)(J) at NMJ6-7 of indicated genotypes. Neto-α is required in motor neurons for normal basal neurotransmission. Scale bars: 1 µm. Error bars indicate SEM. ****, *p*<0.0001; ***, *p*<0.001; ns, p>0.05. Genotypes: control (*w^1118^*), muscle rescue (*neto-α^null^;G14-Gal4/UAS-neto-α),* neuron rescue (*neto-α^null^;OK6-Gal4/UAS-neto-α* and *neto-α^null^,BG380-Gal4/Y; UAS-neto-α/+),* muscle RNAi *(G14-Gal4/+; UAS-neto-α^RNAi^/+*), neuron RNAi *(BG380-Gal4/+;; UAS-neto-α^RNAi^/+*).

Surprisingly, muscle overexpression of *neto-α* could not rescue the neurotransmission defects of *neto-α^null^* mutants (Figures 4H-J, Supplemental Table 1). The EJPs amplitude and quantal content remained severely reduced in these animals (EJP, 16.84 ± 1.59 mV and QC, 17.71 ± 1.19). However, neuronal expression of *neto-α* restored all these parameters to control levels (*BG380* rescue EJP, 36.33 ± 1.90 mV and QC, 32.74 ± 2.11). We confirmed these results with multiple motor neuron specific promoters (*BG380-Gal4* and *OK6-Gal4* shown in Figure 4H-J). Furthermore, knockdown of Neto-α in neurons but not in muscles recapitulated the electrophysiological phenotypes of *neto-α^null^* mutants (*BG380>neto-α^RNAi^* EJP, 15.45 ± 1.51 mV and QC, 14.15 ± 1.13; *G14>neto-α^RNAi^* EJP, 31.73 ± 2.22 mV and QC, 24.83 ± 3.40).

Together these data suggest that Neto-α functions in both motor neurons and muscles. In muscles, Neto-α limits the PSD size, whereas in motor neurons, Neto-α has critical roles in ensuring normal neurotransmitter release. These functions and the low endogenous level of Neto-α are in sharp contrast to those of Neto-β, the predominant isoform at larval NMJ. Unlike Neto-α, Neto-β is required for the synaptic recruitment and stabilization of glutamate receptors (Ramos et al., 2015). The *neto-β^null^* NMJs have greatly diminished postsynaptic receptors and thus reduced minis (quantal size) but have normal basal neurotransmission due to a compensatory increase in quantal content. In contrast, both basal neurotransmission and quantal content are diminished in the absence of Neto-α suggesting homeostasis deficits.

### Loss of homeostatic plasticity at *neto-α^null^* NMJs

We tested for a role for Neto-α in the homeostatic control of synaptic function using well-studied chronic and acute homeostasis paradigms (Frank et al., 2006). Deletion of *GluRIIA* subunit greatly diminishes the quantal size throughout NMJ development; this triggers increased quantal content which restores the evoked muscle responses to normal levels (DiAntonio et al., 1999). In our hands, the *GluRIIA^null^* mutants had mEJPs reduced by 50 % (*IIA^null^*, 0.59 ± 0.04 mV vs. *w^1114^*, 1.19 ± 0.05 mV*, p =* 0.0001), quantal content increased by 60 % (*IIA^null^*, 45.70 ± 3.15, compared to 28.91 ± 2.48 in control, *p* = 0.0006), and relatively normal EJP amplitude (*IIA^null^*, 26.80 ± 2.73 mV, compared to 33.79 ± 2.34 mV in control, *p* = 0.07) (Figure 5A-B). This presynaptic compensatory response did not occur in the absence of Neto-α; the EJP amplitude was reduced in *neto-α^null^*; *GluRIIA^null^* double mutants at levels lower than each individual mutant (*neto-α^null^*; *IIA^null^*, 8.02 ± 0.73 mV, compared to *IIA^null^*, 26.80 ± 2.73 mV and *neto-α^null^*, 17.52 ± 2.06 mV, *p* = 0.0001). These double mutants have reduced mEJPs (*neto-α^null^*; *IIA^null^*, 0.50 ± 0.02 mV, compared to *IIA^null^*, 0.59 ± 0.04 mV and *neto-α^null^*, 1.03 ± 0.10 mV, *p* = 0.07) but lack any homeostatic increase in quantal content (*neto-α^null^*; *IIA^null^*, 16.26 ± 1.60, compared to *IIA^null^*, 45.70 ± 3.15 and *neto-α^null^*, 18.09 ± 2.57, *p* < 0.0001). These results are reminiscent of a previously described hypomorphic allele of *neto* (*neto^109^*) with severe deficits in homeostatic plasticity (Kim et al., 2012).

**Figure 5:**
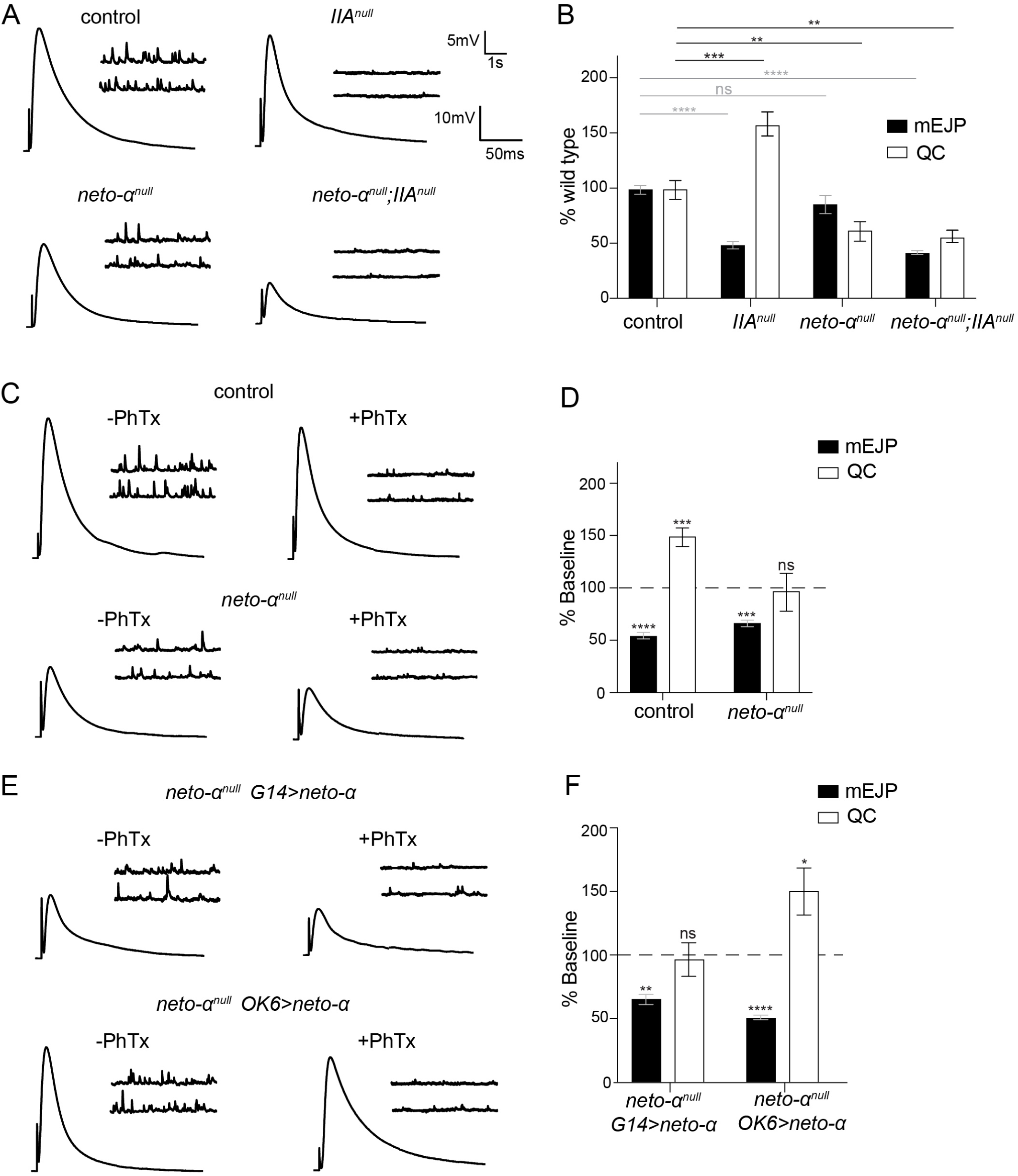
Neto-α is required for the presynaptic homeostatic response. (A) Representative traces for mEJPs and EJPs recordings at 0.5 mM extracellular Ca^2+^ from muscle 6 of indicated genotypes. Note the further reduced EJPs amplitude in *neto-α^null^;IIA^null^* double mutants. (B) Quantification of mEJPs amplitude and QC values normalized to control _(*w*_*_1118_*_)._ (C) Representative traces for mEJPs and EJPs recordings before and after PhTx treatment in control and *neto-α^null^* mutants at 0.5 mM extracellular Ca^2+^. (D) Quantification of mEJPs amplitude and QC values after PhTx treatment and recordings at 0.5 mM extracellular Ca^2+^, normalized to the baseline values of the same genotype. Following PhTx application, *neto-α^null^* mutants fail to restore their basal neurotransmission and show no presynaptic compensatory response (no increase in QC). (F) Representative traces for mEJPs and EJPs recordings before and after PhTx application in *neto-α^null^* mutants rescued by muscle or neuron expressed *neto-α*. (G) Quantification of mEJPs amplitude and QC relative values (after/before PhTx treatment, within the same genotype), show a strong increase in presynaptic release (QC) only in neuronally rescued mutants. Data are represented as mean ± SEM.****, *p*<0.0001; ***, *p*<0.001; **, p<0.05; *, p<0.01; ns, p>0.05.

To examine the speed of the Neto-α-mediated homeostatic response, we used an acute homeostasis paradigm that utilizes Philanthotoxin-343 (PhTx), an effective glutamate receptor blocker (Frank et al., 2006). PhTx applications to dissected NMJ preparations trigger significant homeostatic compensation within ten minutes. Indeed, in control NMJ preparations exposed to 20 μM PhTx, we observed a strong decrease of mEJP amplitude (from 1.25 ± 0.05 mV to 0.69 ± 0.04 mV) and a robust compensatory response, with quantal content increasing from 31.31 ± 2.40 to 47.13 ± 2.77 (Figure 5C-D). PhTx applications also triggered reduced mEJP at *neto-α^null^* NMJs (from 1.09 ± 0.06 mV to 0.74 ± 0.03 mV); however, *neto-α^null^* did not show any changes in quantal content (from 19.75 ± 1.63 to 19.37 ± 3.58). Similar recordings performed at higher Ca^2+^ concentration (0.8 mM Ca^2+^) showed increased EJP amplitude at both control and *neto-α^null^* NMJs (56.93 ± 2.44 mV in control and 67.08 ± 4.40 mV in *neto-α^null^*) (Figure 5E). However, no substantial compensatory response/ increase in quantal content was observed in the absence of Neto-α (66.95 ± 5.48 before and 73.15 ± 6.26 after PhTx). These results demonstrate that Neto-α is critical for both chronic and acute homeostatic modulation of neurotransmitter release in response to reduced postsynaptic sensitivity.

We next tested the tissue specific requirements for Neto-α in homeostatic plasticity using rescue experiments and acute PhTx applications on NMJ preparations. Overexpression of *neto-α* in motor neurons, but not in muscles, significantly rescued the PhTx-induced increase in quantal content at *neto-α^null^* NMJs (to 48.97 ± 5.60 in *neto-α^null^*; *OK6>neto-α* vs. 17.31 ± 2.35 in *neto-α^null^*; *G14*>*neto-α*) (Figure 5F-G). Together these results demonstrate that Neto-α functions in the motor neurons to modulate basal neurotransmission and to confer robust homeostatic plasticity. Since rapid homeostatic compensation occurs in NMJ preparations with severed motor axons, in the absence of either protein translation or action potential-induced evoked neurotransmission (Frank et al., 2006), these results suggest that Neto-α functions in the presynaptic terminals or is developmentally required for a presynaptic activity required for PHP.

Using an antisense probe specific to the Neto-α intracellular domain, we found that *neto-α* transcript is expressed in the striated muscle starting from late embryo stages through larval stages (third instar control and *neto-α^null^* shown in Figure 6A-B). Long exposure also revealed *neto-α* expression in a subset of cells in the larval central nervous system. Overexpression of tagged Neto-α in motor neurons produced accumulation of Neto-α-positive signals at synaptic terminals, along the axons, and in the somato-dendritic compartment of these neurons within the ventral ganglia (Figure 6C). Regardless of the nature of the tag (GFP or V5-not shown), these signals appeared as distinct puncta in the axonal compartment suggesting that Neto-α either concentrates in secretory vesicles or forms aggregates at the neuron surface. Intriguingly, neuronal overexpression of Neto-β did not induce accumulation along axons or at the synaptic terminals; instead, Neto-β remained restricted to the somato-dendritic compartment (Figure 6D, see below).

**Figure 6:**
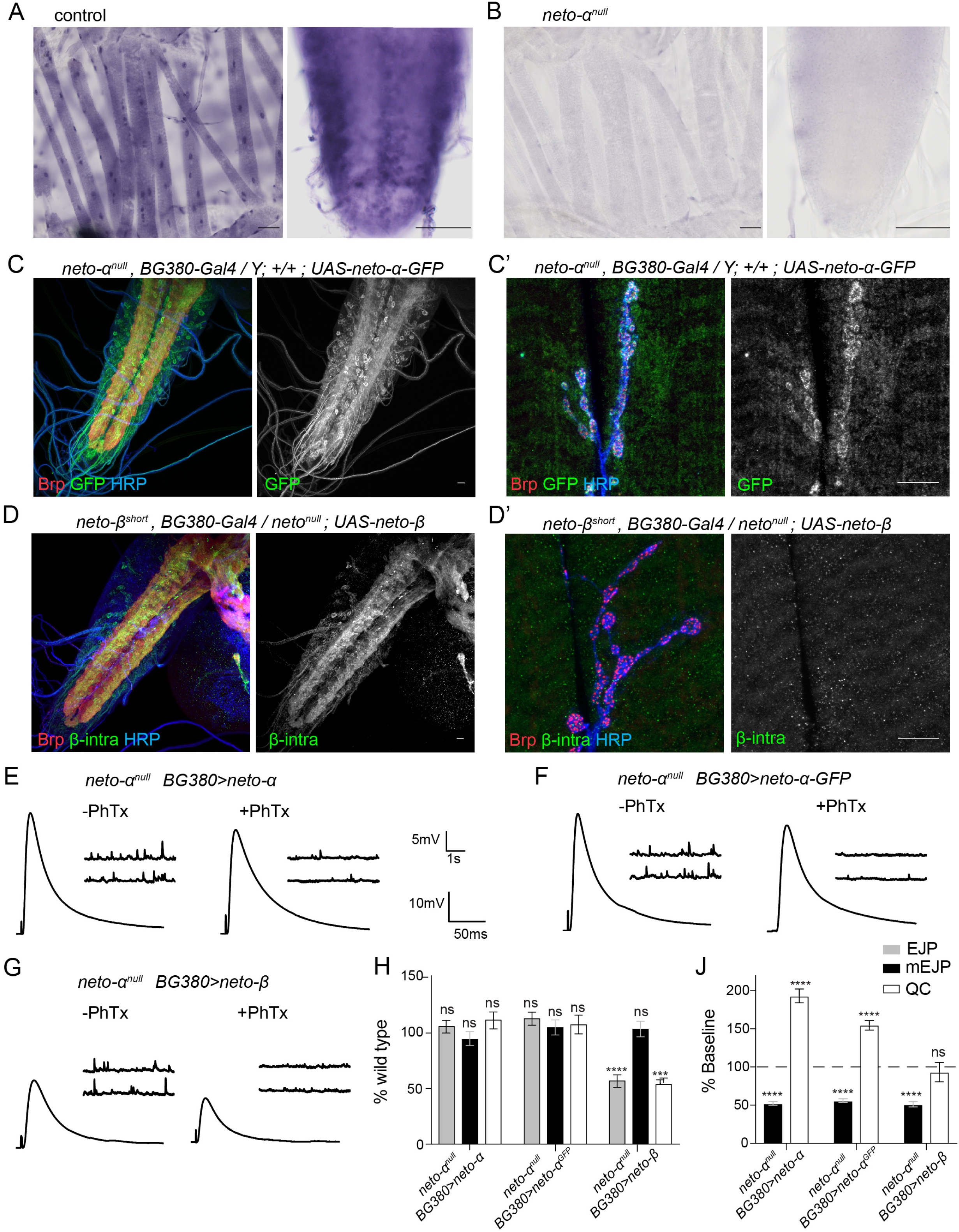
Different distribution and function for Neto-α and Neto-β. (A-B) Expression of *neto-α* specific exon in the striated muscles and ventral ganglia of third instar larvae by in situ hybridization. (C-D) Confocal images of the ventral ganglia (C-D) and NMJ boutons (C’-D’) labeled for Brp (red), GFP or Neto-β-intra (green) and HRP (blue) showing the distribution of Neto-α-GFP (C) and Neto-β (D) when overexpressed in motor neurons. Neto-α-GFP labels the motor neuron soma and axons, and accumulates in a punctate pattern at synaptic terminals, even in the absence of endogenous Neto-α. In contrast, Neto-β does not label the axons and could not be detected at synaptic terminals. Note that anti-β-intra antibodies recognize a C-terminal peptide that is missing in *neto-β ^short^*, thus facilitating unambiguous detection of full length Neto-β. (E-G) Representative traces for mEJPs and EJPs recordings for the indicated genotypes before and after PhTx treatment. (H) Quantification of mEJPs amplitude, EJP amplitude, and QC values normalized to control (*w^1118^*). (J) Quantification of mEJPs amplitude and QC relative values (after/before PhTx treatment, within the same genotype). Unlike *neto-α* and *neto-α-GFP*, *neto-β* overexpression in motor neurons cannot rescue the electrophysiological and homeostasis deficits of *neto-α^null^* mutants. Scale bars: 50 µm (A-B), 10 µm (C-D). Data are represented as mean ± SEM.****, *p*<0.0001; ***, *p*<0.001; ns, p>0.05.

The tags do not interfere with the presynaptic functions of Neto-α: Tagged Neto-α variants rescued the basal neurotransmission of *neto-α^null^* mutants as effectively as unmodified Neto-α (GFP-tagged vs. no tag shown in Figure 6E-F). Also, expression of Neto-α-GFP in the motor neurons was similar to Neto-α in restoring the acute homeostatic response at *neto-α^null^* NMJs (quantified in Figure 6H-J). More specifically, neuronal expression of *neto-α* or *neto-α-GFP* rescued the EJP amplitude in *neto-α^null^* mutants to 36.33 ± 1.90 mV and 38.69 ± 1.97 mV, respectively; in response to PhTx application, the quantal content increased from 32.74 ± 2.12 to 63.30 ± 2.93 in *neto-α* rescued mutants, and from 31.54 ± 2.66 to 49.99 ± 1.81 in *neto-α-GFP* rescued animals. In contrast, expression of *neto-β* in motor neurons could not rescue basal neurotransmission or homeostatic potentiation at *neto-α^null^* NMJs (Figure 6G-H). These larvae exhibit reduced basal neurotransmission (19.73 ± 1.89 mV before and 9.39 ± 1.25 after PhTx application) and no compensatory increase in quantal content (16.00 ± 1.20 vs 14.97 ± 2.02), resembling the *neto-α^null^* NMJs. Lack of any Neto-β-mediated neuronal rescue could reflect the inability of Neto-β to localize to presynaptic terminals, and/or to fulfill the Neto-α specific functions in motor neurons. These results uncover new isoform specific functions for Neto-α at the *Drosophila* NMJ and suggest that Neto-α function in the presynaptic terminal.

### Neto-α enables the fast recruitment of the active zone protein Brp

The neurotransmission defects at *neto-α^null^* NMJs may reflect deficits in presynaptic Ca^2+^ entry. We investigated this possibility using a Ca^2+^-sensitive fluorescent dye loaded into motor nerve terminals (Macleod, 2012). This dye fluoresces in proportion to free Ca^2+^ levels in the cytosol ([Ca^2+^]_c_); when loaded in constant proportion to a Ca^2+^ insensitive dye, it allows ratiometric comparisons between *neto-α^null^* and control type-Ib and -Is terminals (Figure 7A). [Ca^2+^]_c_ at rest, estimated prior to stimulation, was no different in *neto-α^null^* relative to control (Figure 7B; Ib: P=0.85; Is: P=0.96). The amplitude and decay of single AP evoked changes in [Ca^2+^]_c_ in response to 1Hz nerve stimulation were no different in *neto-α^null^* (amplitude, Figure 7C-D: Ib: P=0.62; Is: P=0.96) (decay: data not shown, Ib: P=0.42; Is: P=0.56). Finally, the Ca^2+^ signals evoked by 10 and 20Hz stimulus trains were no different in *neto-α^null^* (10Hz, data not shown: Ib: P=0.63; Is: P=0.93) (20Hz, Figure 7E: Ib: P=0.37; Is: P=0.40). While the data shown here are sufficiently sensitive to reveal the differences in Ca^2+^ entry known to exist between type-Ib and -Is terminals (He et al., 2009; Lu et al., 2016), they reveal no deficit in Ca^2+^ entry in *neto-α^null^* terminals. Thus, *neto-α^null^* neurotransmission deficits are most likely the result of deficits in the release apparatus downstream of Ca^2+^ entry.

**Figure 7:**
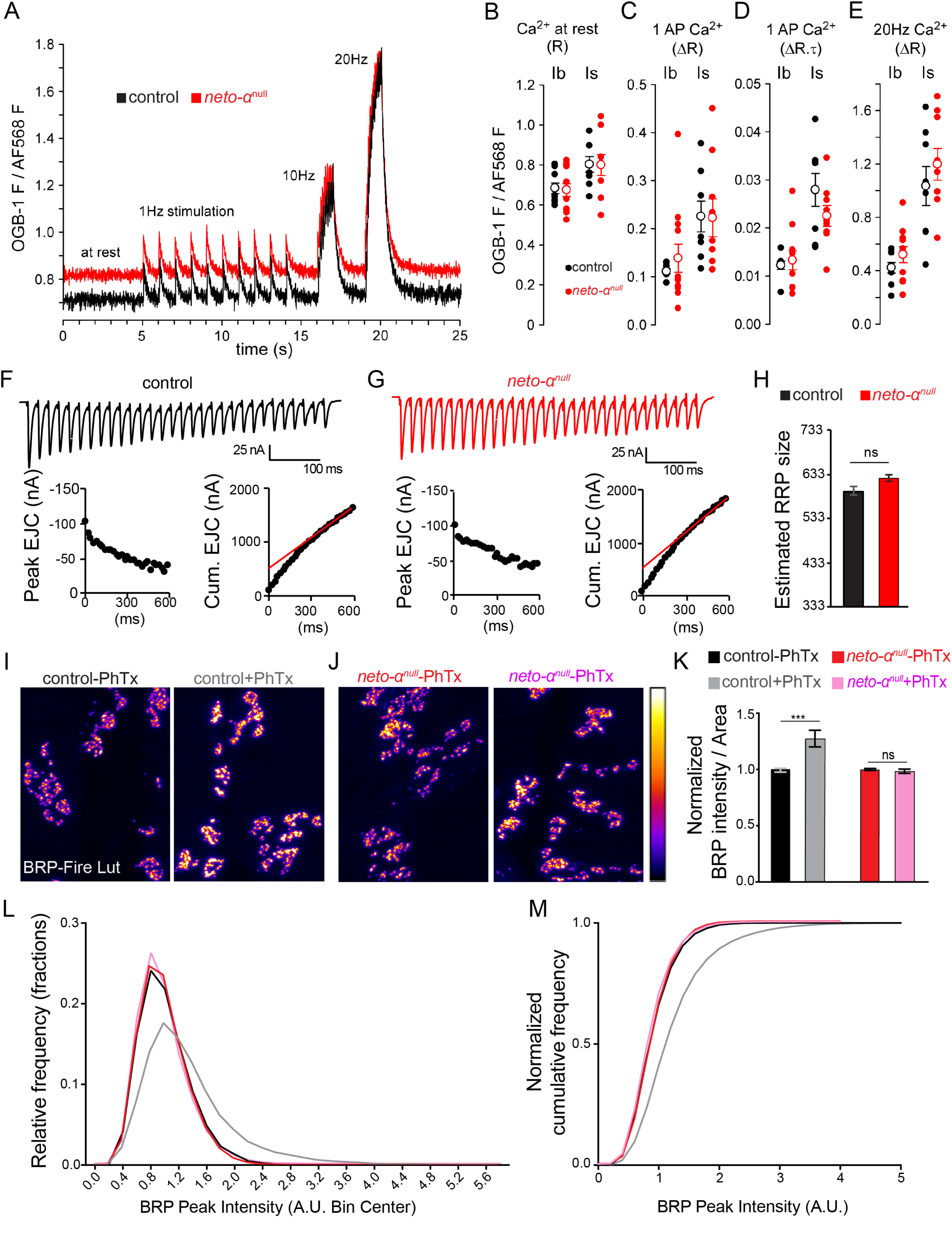
Cytomatrix remodeling is impaired at *neto-α^null^* NMJs. (A-E) Action-potential mediated Ca^2+^ transients are no different in *neto-α^null^* motor neuron terminals relative to control. (A) Single trial traces of changes in Ca^2+^-sensitive Oregon Green BAPTA-1 (OGB-1) fluorescence relative to Ca^2+^-insensitive Alexa Fluor 568 (AF568) fluorescence in the cytosol of a type-Ib terminals on muscle 6, in response to stimuli applied to the hemisegment nerve: 10@1Hz, 10@10Hz and 20@20Hz. OGB-1 images collected at 112 frames per second. (B) Scatter plot of OGB-1 / AF568 fluorescence prior to nerve stimulation, representing free Ca^2+^ levels in the cytosol ([Ca^2+^]_c_) at rest. Each closed circle represents a ratio (R) measurement from a specific terminal type (Ib or Is) in a different larva. Open circles represent mean ± SEM. (C) Scatter plot of the amplitude (change in ratio: ΔR) of Ca^2+^ transients evoked by stimuli delivered at 1Hz. (D) Scatter plot of the product of amplitude (ΔR) and decay time course (τ; reported in seconds) of Ca^2+^ transients evoked by 1Hz stimuli (ΔR. τ). (E) Scatter plot of the amplitude (change in ratio: ΔR) of Ca^2+^ transients evoked by a 10Hz train of stimuli. All data collected from muscle 6, segment A4, in 0.5 mM Ca^2+^ HL3. P values from Student’s T-tests are reported in the text. The Mann Whitney U Test was applied when normality tests failed. (F-H) Representative EJC traces (top) and cumulated peak EJC amplitudes (bottom) for 30 stimuli at 50 Hz at 1.5 mM extracellular Ca^2+^ in control (F) and *neto-α^null^* (G). (H) Estimated RRP sizes for control and *neto-α^null^* are similar (n =5, *p* = 0.1441). (I-J) Quantification of BRP intensity following 10 minute of vehicle or PhTx treatment at control and *neto-α^null^* NMJs. BRP is showed in Fire-lut; on the intensity scale, white represents peak intensity (20,000 arbitrary units, A.U.). PhTx application induces a 27.50 ± 0.07% increase in Brp signal intensity in control (n=24 NMJs without and 26 with PhTx, *p*=0.0008), but not in *neto-α^null^* mutants (n=19 and 22, *p*=0.4941). (K-L) Normalized frequency distribution (K) and cumulative frequency (L) of BRP peak intensities reveal a rightward shift after PhTx application for the control animals (from 1 to 1.34, n=18312 peaks without and 18756 with PhTx, *p*<0.0001), but not for *neto-α^null^* (from 1 to 0.97, n=14063 without and 17940 with PhTx, *p*<0.0001). Data are represented as mean ± SEM. ****, *p*<0.0001; ***, *p*<0.001; ns, p>0.05.

To estimate the number of release-ready presynaptic vesicles in *neto-α^null^* mutants, we analyzed cumulative postsynaptic current during high-frequency stimulus trains (30 stimuli at 50 Hz) as previously described (Muller et al., 2012) (Figure 7F-H). Briefly, we measured evoked excitatory junction currents (EJCs) at a voltage clamped to – 65 mV, in HL-3 saline with 1.5 mM Ca^2+^ and 10 mM Mg^2+^, and cumulated EJCs evoked by 50 Hz stimulation (30 stimuli) of control and *neto-α^null^* NMJs (Materials and Methods). Back-extrapolation from linear fits to the cumulative EJC to time zero yielded 418 ± 22 vesicles for control and 451 ± 18 for *neto-α^null^* (n= 5, *p* = 0.28). Finally, the size of the RRP pool was calculated, and there was no significant difference between control (522 ± 28, n = 5) and *neto-α^null^*, (581 ± 24, n =5, *p* = 0.14). This result indicates that the absence of Neto-α does not alter the basal RRP size and therefore could not cause the observed reduced basal neurotransmission.

Previous studies demonstrated that PhTx application results in a rapid increase in the quantity of presynaptic active zone protein Brp, accompanied by an elaboration of the presynaptic cytomatrix structure (Goel et al., 2017; Weyhersmuller et al., 2011). We visualized and quantified the Brp puncta before and after PhTx exposure by confocal microscopy (see Materials and Methods). As expected, upon PhTx application, control NMJs showed a significant increase (27.50 ± 0.07%) in Brp-positive immunoreactivities (n=24 NMJs without and 26 with PhTx, *p*=0.0008) (Figure 7I-J). However, no increase in the Brp-positive signals was detectable at *neto-α^null^* NMJs. Furthermore, relative frequency and cumulative probability distributions of Brp intensities revealed a rightward shift only in PhTx-treated control but not *neto-α^null^* NMJs (Figures 7K-L). These findings suggest that, in response to PhTx-triggered reduced postsynaptic sensitivity, Neto-α functions to swiftly mobilize the active zone protein Brp, which presumably enhances vesicle release and enables the compensatory response.

### Distinct domains of Neto-α regulate basal release and presynaptic potentiation

We have previously demonstrated that a minimal Neto variant, called Neto-ΔCTD (including the highly conserved extracellular CUB domains, LDLa motif, the transmembrane part, but no intracellular C-terminal domain) is both required and sufficient for the synaptic recruitment and function of postsynaptic KARs (Ramos et al., 2015). Neuronal overexpression of Neto-ΔCTD-GFP recapitulated the distribution of Neto-α-GFP, and localized to dendrites and soma, along axons, and at synaptic terminals (Figure 8A, compare with Figure 6C). Importantly, neuronal Neto-ΔCTD-GFP rescued the basal neurotransmission at *neto-α^null^* mutant NMJs (EJP, 37.15 ± 0.98 mV and QC, 49.99 ± 1.81, Figure 8C-C’). This indicates that Neto-ΔCTD is sufficient for normal basal neurotransmission. Note that these animals have decreased mEJPs amplitude and a slightly increased QC, suggesting some compensatory developmental response (Figure 8C’). However, this variant could not rescue the acute PHP response in *neto-α^null^* mutants (Figure 8C”). Instead, upon PhTx exposure the quantal content decreased from 39.15 ± 1.67 to 23.47 ± 2.78 at these NMJs, indicating that the intracellular part of Neto-α, albeit dispensable for basal neurotransmission, is absolutely required for PHP.

**Figure 8:**
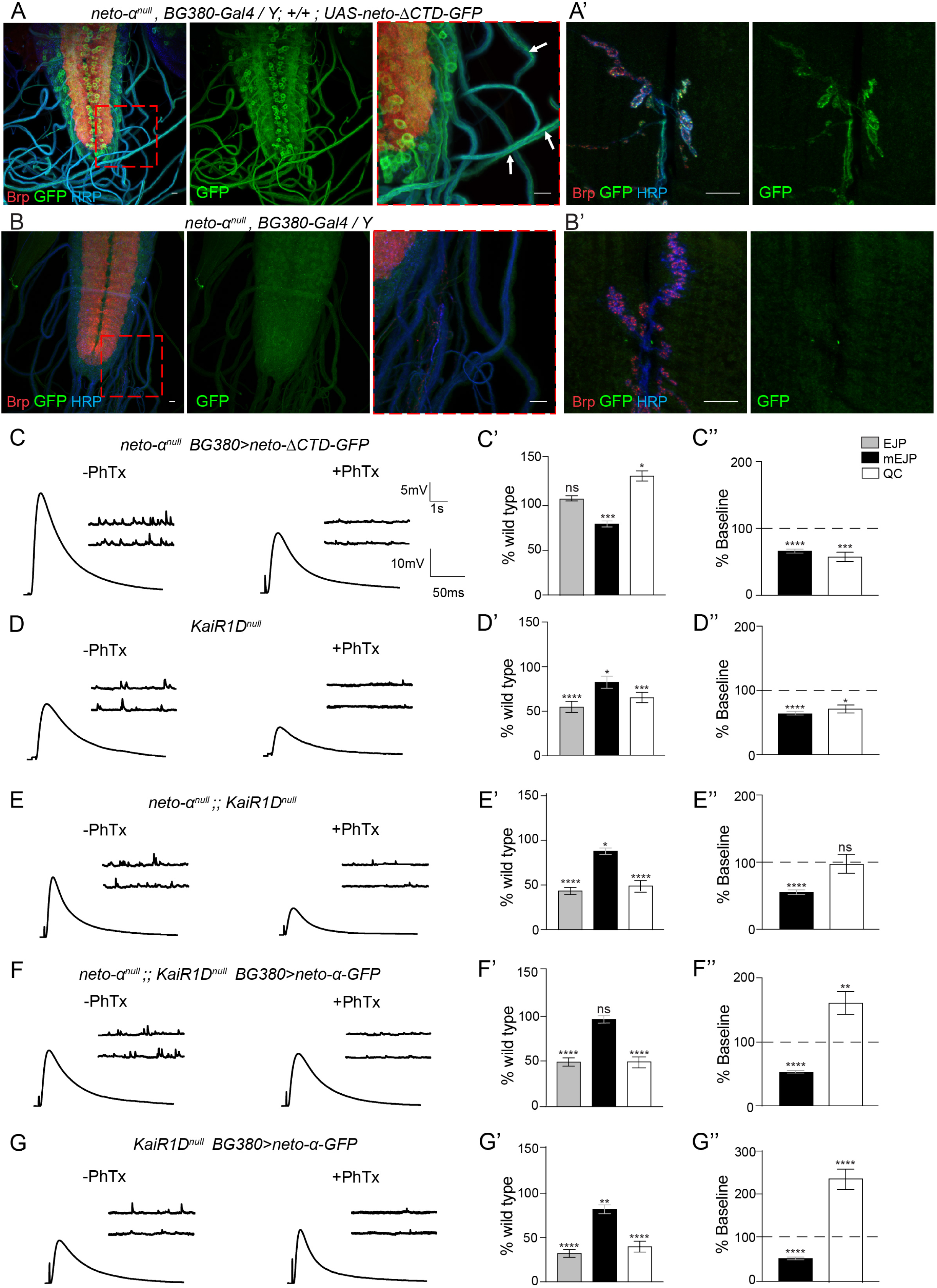
Distinct domains of Neto-α regulate basal release and presynaptic potentiation. (A-B) Confocal images of the ventral ganglia and NMJ boutons (A’-B’) labeled for Brp (red), GFP (green) and HRP (blue) showing the distribution of Neto-ΔCTD-GFP when overexpressed in motor neurons (A) and the negative control (B). Similar to Neto-α-GFP (Figure 6C), Neto-ΔCTD-GFP labels the soma and axons of motor neurons and accumulates at synaptic terminals. Note that neuronal Neto-ΔCTD-GFP can rescue the basal neurotransmission defects at *neto-α^null^* NMJs but cannot restore the homeostatic response (below). (C-G) Sets of electrophysiological recordings of basal neurotransmission and presynaptic homeostatic potentiation response for the indicated genotypes. Each analysis includes representative traces for mEJPs and EJPs recordings before and after PhTx application (left), quantification of mEJPs amplitude, EJP amplitude, and QC values normalized to control (*w^1118^*) (middle), and quantification of mEJPs amplitude and QC relative values after PhTx treatment, normalized to the baseline values of the same genotype. The electrophysiological defects of *KaiRID^null^* NMJs resemble those of *neto-α^null^* mutants, as well as of *neto-α^null^;;KaiRID^null^* suggesting that KaiRID and Neto-α function in the same pathway (D-E). Overexpression of *neto-α-GFP* in *neto-α^null^;;KaiRID^null^* motor neurons does not alleviate the ∼50% reduction in EJP amplitude, but enables a significant PHP response (F). When *neto-α-GFP* is overexpressed in the presence of endogenous Neto-α, the amplitude of the PHP response is dramatically increased (G). Scale bars: 10 µm. Data are represented as mean ± SEM. ****, *p*<0.0001; ***, *p*<0.001; ns, p>0.05.

Recent studies have described a presynaptic KAR subunit, KaiRID, which controls basal neurotransmission and confers PHP at the fly NMJ (Kiragasi et al., 2017; Li et al., 2016). Since Neto proteins modulate the function of KARs, the phenotypic similarities between *neto-α^null^* and *KaiRID* loss-of-function mutants suggest that Neto-α partly functions by modulating the presynaptic KaiRID. In both mutants, neuronal expression of the corresponding full-length transgenes rescued the basal neurotransmission and PHP deficits (KaiRID loss-of-function: (Kiragasi et al., 2017) and *neto-α^null^*: Figure 5). However, a Ca^2+^-impermeable variant (*KaiRID^R^)* restored the presynaptic homeostasis at *KaiRID* mutant NMJs, but could not rescue the basal neurotransmission (Kiragasi et al., 2017; Li et al., 2016). Since presynaptic Neto-ΔCTD efficiently rescued the EJP amplitudes at *neto-α^null^* NMJs, but only full-length Neto-α could rescue their PHP, Neto-α may (i) engage KaiRID and modulate basal neurotransmission via KaiRID/Neto-ΔCTD complexes, and (ii) confer homeostatic potentiation via its intracellular part.

We tested this model by first generating a *KaiRID^null^* mutant using the CRISPR/Cas9 methodology and comparing the phenotypes of single and double (*neto-α^null^* and *KaiRID^null^*) mutants (Methods and Figure 8D-E). The *KaiRID^null^* neurotransmission defects were fully rescued by expression of KaiRID in the motor neurons, confirming the specificity of the molecular lesion (Supplemental Table 1). The basal neurotransmission defects observed at *KaiRID^null^* synapses recapitulated the phenotypes reported for *KaiRID* loss-of-function alleles (Kiragasi et al., 2017), and were very similar to those observed for *neto-α^null^* mutants (18.80 ± 2.09 mV for *KaiRID^null^* vs 17.52 ± 2.06 mV for *neto-α^null^*) (Figure 8D, compare with Figure 5C-D). The double mutant showed basal neurotransmission defects within the range of single individual mutants (15.49 ± 1.41 mV) (Figure 8E). These results suggest that Neto-α and KaiRID function together in motor neurons to control basal neurotransmission. *neto-α^null^* and *KaiRID^null^* single and double mutants were also similarly impaired in their acute PHP responses (Figures 8D-E and Supplemental Table 1). Since the intracellular part of Neto-α is key to PHP, we examined whether expression of Neto-α-GFP in motor neurons could partly rescue the neurotransmission defects at *neto-α^null^;;KaiRID^null^* NMJs. As expected, neuronal expression of Neto-α-GFP did not rescue the basal neurotransmission at *neto-α^null^;;KaiRID^null^* NMJs, which remained at 17.45 ± 1.56 mV (Figure 8F). However, these animals showed normal homeostatic response; their quantal content was 15.03 ± 1.56 before PhTx and 24.63 ± 2.67 after PhTx. Interestingly, neuronal overexpression of Neto-α in the *KaiRID^null^* single mutant background exacerbated the amplitude of the PHP response; the quantal content changed from 11.95 ± 1.83 before PhTx to 29.31 ± 2.81 after PhTx, (Figure 8G). This indicates that (i) Neto-α is sufficient for presynaptic homeostasis, and (ii) endogenous levels of Neto-α are limiting. Overexpression of Neto-ΔCTD in the *KaiRID^null^* background did not restore basal neurotransmission or PHP (not shown) due to missing key determinants: KaiRID for basal neurotransmission, and Neto-α intracellular part for homeostasis. Together, these data demonstrate that the two major functions of Neto-α in the presynaptic compartment could be segregated and mapped to different domains: 1) the minimal Neto, Neto-ΔCTD, which modulates basal neurotransmission, likely by modulating the KaiRID function, and 2) the intracellular part of Neto-α, which is both required and sufficient for the presynaptic homeostatic response.

## Discussion

Here we show that Neto-α is required in both pre- and post-synaptic compartments for the proper organization and function of the *Drosophila* NMJ. The fly NMJ is a glutamatergic synapse that utilizes at least six distinct KAR subunits; they form two distinct postsynaptic complexes (type-A and type-B) that co-exist within individual PSDs and enable NMJ functionality and plasticity, and a presynaptic KaiRID-containing complex that modulates basal neurotransmission. In muscle, Neto-α limits the size of the postsynaptic receptors field; the PSDs are significantly enlarged in muscle where Neto-α has been perturbed (Figures 3-4). In motor neurons, Neto-α is required for two distinct activities: (1) modulation of basal neurotransmission in a KaiRID-dependent manner, and (2) effector of presynaptic homeostasis response. To our knowledge this is the first example of a glutamate receptors auxiliary protein that modulates receptors on both sides of a particular synapse and plays a distinct role in homeostatic plasticity.

Vertebrate KARs depend on Neto proteins for their distribution and function (Copits and Swanson, 2012). Due to their reliance on KARs, *Drosophila neto^null^* mutants have no functional NMJs (no postsynaptic KARs) and consequently die as completely paralyzed embryos (Kim et al., 2012). We have previously shown that muscle expression of Neto-ΔCTD, or “minimal Neto”, rescues, at least in part, the recruitment and function of KARs at synaptic locations (Han et al., 2015; Kim et al., 2012; Kim et al., 2015; Ramos et al., 2015). Here we report that neuronal Neto-ΔCTD also rescues the KaiRID-dependent basal neurotransmission (Figure 8). Thus Neto-ΔCTD, the highly conserved segment of Neto, seems to represent the Neto core of KAR modulatory activities.

The intracellular parts of Neto proteins are highly divergent, likely reflecting the microenvironments where different Neto proteins operate (Copits and Swanson, 2012; Tomita, 2010). Similar to mammalian Neto1 and Neto2, *Drosophila* Neto-α and Neto-β are differentially expressed in the CNS (not shown) and have different intracellular domains that mediate distinct functions. These large intracellular domains are rich in putative phosphorylation sites and docking motifs and could further modulate the distribution and function of KARs or serve as signaling hubs and protein scaffolds. Post-translational modifications regulate vertebrate Neto activities *in vitro,* albeit the *in vivo* relevance of these changes remains unknown (Lomash et al., 2017). Our data demonstrate that Neto-α and Neto-β could not substitute for each other (this study and (Ramos et al., 2015)). For example, Neto-β, but not Neto-α, controls the recruitment of PAK, a PSD component that stabilizes selective KARs subtypes at the NMJ, and ensures proper postsynaptic differentiation (Ramos et al., 2015). Conversely, postsynaptic Neto-β alone cannot maintain a compact PSD size; muscle Neto-α is required for this function (Figure 4). Importantly, Neto-β cannot fulfill any of the presynaptic functions of Neto-α, presumably because is confined to the somato-dendritic compartment and cannot reach the synaptic terminals (Figure 6D). Histology and Western blot analyses indicate that Neto-α constitutes less than 1/10 of the net Neto at the *Drosophila* NMJ (Figure 1 and (Ramos et al., 2015)). These low levels impaired our ability to directly visualize endogenous Neto-α. We have generated several isoform specific antibodies but they could only detect Neto-α when overexpressed (not shown). Similar challenges have been encountered in the vertebrate Neto field (Wyeth et al., 2017).

Interestingly, the two Neto isoforms are limiting in different synaptic compartments. Neto-β limits the recruitment and synaptic stabilization of postsynaptic KARs (Ramos et al., 2015). In contrast, several lines of evidence indicate that Neto-α is limiting in the motor neurons: First, overexpression of KaiRID cannot increase basal neurotransmission (Kiragasi et al., 2017), but strong neuronal overexpression of Neto-ΔCTD increases the basal neurotransmission (not shown), indicating that Neto and not KaiRID is limiting in the motor neurons. Secondly, neuronal overexpression of Neto-α exacerbates the PHP response to PhTx exposure and even rescues this response in *KaiRID^null^* (Figure 8). These findings suggest that KaiRID’s function during PHP is to help traffic and stabilize Neto-α, a low abundant PHP effector. Similarly, studies in mammals reported that kainate receptors trafficking in the CNS does not require Neto proteins, but rather kainate receptors regulate the surface expression and stabilization of Neto1 and −2 (Straub et al., 2011; Zhang et al., 2009). Nonetheless, the KAR-mediated stabilization of Neto proteins at CNS synapses supports KARs distribution and function. In flies, the KaiRID-dependent Neto-α delivery at synaptic terminals ensures both a KAR-dependent function, normal basal neurotransmission, and a Neto-α-specific activity as an effector of PHP.

Previous studies showed that presynaptic KARs function to regulate neurotransmitter release; however, the site and mechanism of action of presynaptic KARs have been difficult to pin down unambiguously (Perrais et al., 2010). This study provides strong evidences for Neto activities at presynaptic terminals. First, Neto-α is both required and sufficient for PHP (Figures 5 and 8). Previous work demonstrated that the fast expression of PHP in response to PhTx occurs even when the motor neuron axon is severed (Frank et al., 2006). Also, the signaling necessary for PHP expression is restricted to postsynaptic densities and presynaptic boutons (Li et al., 2018). Second, Neto-ΔCTD, but not Neto-β, rescued the basal neurotransmission defects in *neto-α^null^* (Figures 6 and 8). Both variants contain the “minimal Neto” required for KARs modulation (Ramos et al., 2015), but only Neto-ΔCTD can reach the presynaptic terminal, while Neto-β is restricted to the somato-dendritic compartment (Figures 6 and 8). This suggest that Neto-ΔCTD (or Neto-α) together with KaiRID localize at presynaptic terminals, where KaiRID could function as an autoreceptor. Finally, upon PhTx exposure, Neto-α enabled the fast recruitment of Brp at the active zone (Figure 7). Multiple homeostasis paradigms trigger Brp mobilization, followed by remodeling of presynaptic cytomatrix (Goel et al., 2017). These localized activities further support Neto-α functioning at presynaptic terminals.

Presynaptic activities for Neto-α include KaiRID modulation [(Li et al., 2016) and this study]. Rapid application of glutamate to outside-out patches from HEK cells transfected with KaiRID indicated that KaiRID forms rapidly desensitizing channels (Li et al., 2016); addition of Neto increases the desensitization rates and open probability for this channel (Han et al, manuscript in preparation). In addition, Neto-α has a large intracellular domain (250 residues) rich in post-translational modification sites and docking motifs, including putative phosphorylation sites for CaMKII, PKC and PKA. This intracellular domain may engage in finely tuned interactions that allow Neto-α to (1) further modulate the KaiRID properties and distribution in response to cellular signals; and (Letts et al.) function as an effector of presynaptic homeostasis in response to low postsynaptic glutamate receptor activity. Mammalian Neto1 and −2 are phosphorylated by multiple kinases *in vitro* (Lomash et al., 2017); CaMKII and PKA-dependent phosphorylation of Neto2 restrict GluK1 targeting to synapses *in vivo* and *in vitro.* Similarly, Neto-α may function in a kinase-dependent manner to stabilize KaiRID and/or other presynaptic components. Secondly, Neto-α may recruit Brp (Figure 7) or other presynaptic molecules that mediate activity-related changes in glutamate release at the fly NMJ. Besides Brp, several presynaptic components have been implicated in the control of PHP (reviewed in (Frank, 2014)). They include (1) Cacophony, the α1 subunit of Ca_V_2-type calcium channels and its auxiliary protein α2δ-3, which control the presynaptic Ca^2+^ influx (Muller and Davis, 2012; Wang et al., 2016), (Letts et al.) the signaling molecules upstream Cac, Eph, Ephexin and Cdc42 (Frank et al., 2009), and (3) the BMP pathway components, Wit and Mad, required for the retrograde BMP signaling (Goold and Davis, 2007). In addition, expression of PHP requires molecules that regulate vesicle release and the RRP size, such as RIM (Muller et al., 2012), Rab3-GAP (Muller et al., 2011), Dysbindin (Dickman and Davis, 2009), SNAP25 and Snapin (Dickman et al., 2012). Recent studies demonstrated that trans-synaptic Semaphorin/Plexin interactions control synaptic scaling in cortical neurons in vertebrates (Wang et al., 2017) but also drive PHP at the fly NMJ (Orr et al., 2017). Neto-α may interact with one or several such presynaptic molecules and function as an effector of PHP. Future studies on what Neto-α cytoplasmic domain binds to and how is it modulated by post-translational modifications should provide key insights into the understanding of molecular mechanisms of homeostatic plasticity.

On the muscle side, Neto-α activities may include (1) engaging scaffolds that limit the PSD size, and (2) modulating postsynaptic KARs distribution and function. For example, Neto-α may recruit trans-synaptic complexes such as Ten-a/Ten-m, or Nrx/Nlgs that have been implicated in limiting the postsynaptic fields (Banovic et al., 2010; Mosca et al., 2012). In particular, DNlg3, like Neto-α, is present in both pre- and postsynaptic compartments and has similar loss-of function phenotypes, including smaller boutons with larger individual PSDs, and reduced EJPs amplitudes (Xing et al., 2014). Neto-α may also indirectly interact with the *Drosophila* PSD95, Dlg, and help establish the PSD boundaries (Figure 3). Fly Netos do not have PDZ-binding domains, but the postsynaptic Neto/KARs complexes contain GluRIIC, a subunit with a class II PDZ-binding domain (Marrus et al., 2004). It has been reported that mutations that change the NMJ receptors gating behavior alter their synaptic trafficking and distribution (Petzoldt et al., 2014). Neto-α could be key to these observations as it may influence both receptor’s gating properties and ability to interact with synapse organizers.

Phylogenetic analyses indicate that Neto-β is the ancestral Neto; in insects, Neto-β is predicted to control the NMJ development and function, including recruitment of iGluRs and PSD components, and postsynaptic differentiation (Ramos et al., 2015). Neto-α appears to be a new, rapidly evolving isoform present in higher *Diptera*. This large order of insects is characterized by a rapid expansion of the KARs branch to ten distinct subunits (Li et al, 2016). Insect KARs have unique ligand binding profiles, strikingly different from vertebrate KARs; however, like vertebrate KARs, they all seem to be modulated by Neto proteins. We speculate that the rapid expansion of KARs forced the diversification of the relevant accessory protein, Neto, and the extension of its repertoire. In flies, the *neto* locus acquired an additional exon and consequently an alternative isoform with distinct expression profiles, subcellular distributions, and isoform specific functions. It will be interesting to investigate how flies differentially regulate the expression and distribution of the two Neto isoforms and control their tissue- and synapse-specific functions. Mammals have five KAR subunits, three of which have multiple splice variants that confer rich regulation (Lerma and Marques, 2013). In addition, mammalian Neto proteins have fairly divergent intracellular parts that presumably further integrate cell specific signals and fine-tune KARs localization and function. In *Diptera*, KARs have relatively short C-tails, and thus limited signaling input, whereas Netos have long cytoplasmic domains that could function as scaffolds and signaling hubs. Consequently, most of the information critical for NMJ assembly and postsynaptic differentiation has been outsourced to the intracellular part of Neto-β (Ramos et al., 2015). Similarly, Neto-α-mediated intracellular interactions may hold key insights into the mechanisms of homeostatic plasticity, as our study reveals that Neto functions as a bona fide effector of presynaptic homeostasis.

### Experimental Procedures

#### Fly stocks

The *neto*-*α^null^* and *KaiRID^null^* alleles were generated using classic CRISPR/Cas9 methodology as previously described (Gratz et al., 2015). Briefly, for each allele, two pairs of gRNAs were injected in *y sc v; [nos-Cas9]attP40/CyO* stock (Ren et al., 2013) followed by germline transformation (Rainbow transgenics). A series of unmarked deletions were isolated and molecularly characterized by PCR from genomic DNA (QuickExtractDNA, Epicentre) and sequencing. Putative genetic null alleles have been isolated and confirmed by sequence analysis; they were subsequently moved in an *w^1118^* background and balanced with markers visible during larval stages. The primers used for gRNAs, PCR and sequencing were as follows:

alpha-1 sense: CTTCGGTTTCTGGGGATAAGATGG

alpha-1 antisense: AAACCCATCTTATCCCCAGAAACC

alpha-3 sense: CTTCGGAATATAATGGAAAAATGA

alpha-3 antisense: AAACTCATTTTTCCATTATATTCC

Neto-F1: AGTCCCTTTACCACTCCATTAGCC

Neto-R1: TTGCGAGTGCTTTTGCCTGC

CG3822-gATD1 sense: CTTCGCATTTTGAATTCGTTCGCGA

CG3822-gATD1 antisense: AAACTCGCGAACGAATTCAAAATGC

CG3822-gATD2 sense: CTTCGACAGCTTCCATGCCGGGAAA

CG3822-gATD2 antisense: AAACTTTCCCGGCATGGAAGCTGTC

CG3822-F1: CAAACCCTTGGAGAAATAGGG

CG3822-R1: CTACGATTGAGGTCCCCTTG.

Neto-F1/R1 are predicted to amplify a 15kb product from control animals and 2kb from *neto*-α*^null^*. Line *#*117 missing 13kb (13,506,327-13,519,803), including the entire alpha-specific exon, was selected as *neto*-*α ^null^*.

CG3822-F1/R1 are predicted to amplify a 994bp product from control animals, and 444bp from *KaiRID^null^*. Line #19 has a truncated message that codes for the first 79 residues of KaiRID, followed by three different amino acids and a stop codon.

Other fly stocks used in this study were as follows: *neto^null^ and neto^hypo^* (Kim et al. 2012); *UAS-neto-α (line A9), UAS-neto-α::GFP (line B4) (Kim et al. 2015)*; *neto-β^null^, neto-β^short^, UAS-neto-β (line NB6), UAS-netoΔCTD (line H6y), neto-α^RNAi^, neto-β^RNAi^* (Ramos et al. 2015), *GluRIIA^SP16^* and *Df(2L)cl^h4^* (Petersen et al. 1997) (from A. DiAntonio, Washington University). The *G14-Gal4*, *BG380-Gal4*, and *OK6-Gal4* were previously described.

### Protein analysis and immunohistochemistry

To analyze muscle proteins, wandering third instar larvae were dissected, and all tissues except for the body wall (muscle and cuticle) were removed. The body walls were mechanically disrupted and lysed in lysis buffer (50 mM Tris-HCl, 150 mM NaCl, 1% Triton X-100, 1% deoxycholate, protease inhibitor cocktail (Roche) for 30 min on ice. The lysates were separated by SDS-PAGE on 4%–12% NuPAGE gels (Invitrogen) and transferred onto PVDF membranes (Millipore). Primary antibodies were used at the following dilutions: rat anti-Neto-ex (Kim et al., 2012), 1:1000; anti-Tubulin (Sigma-Aldrich),1:1000.

For immunohistochemistry, wandering third instar larvae of the desired genotypes were dissected in ice-cooled Ca^2+^-free HL-3 solution (70 mM NaCl, 5 mM KCl, 20 mM MgCl_2_, 10 mM NaHCO_3_, 5 mM trehalose, 5 mM HEPES, 115 mM sucrose) (Stewart et al. 1994) (Budnik, Gorczyca, and Prokop 2006). The samples were fixed in 4% paraformaldehyde (PFA) (Polysciences, Inc.) for 20 min or in Bouin’s fixative (Bio-Rad) for 3 min and washed in PBS containing 0.5% Triton X-100. For PhTx treatment, thirds instar larvae were pinned anteriorly and posteriorly, dissected along the dorsal midline and incubated either with 10 µM PhTx for 15 min in Ca^2+^-free HL-3, or without PhTx for the control. PhTx was then washed out, the fat body and guts were removed and the fillets were fixed in 4% PFA for 20 min, then processed normally.

Primary antibodies from Developmental Studies Hybridoma Bank were used at the following dilutions: mouse anti-GluRIIA (MH2B), 1:100; mouse anti-Dlg (4F3), 1:1000; mouse anti-Brp (Nc82), 1:200. Other primary antibodies were utilized as follow: rat anti-Neto-ex, 1:1000 (Kim et al., 2012); rabbit anti-Neto-β, 1:1,000, rabbit anti-GluRIIC, 1:2,000, rabbit anti-GluRIIB, 1:1,000, (Ramos et al. 2015); chicken anti-GFP, 1:1,000, (Abcam); and Cy5- conjugated goat anti-HRP, 1:1000 (Jackson ImmunoResearch Laboratories, Inc.). Alexa Fluor 488-, Alexa Fluor 568-, and Alexa Fluor 647-conjugated secondary antibodies (Molecular Probes) were used at 1:200. All samples were mounted in ProLong Gold (Invitrogen).

Samples of different genotypes were processed simultaneously and imaged under identical confocal settings in the same imaging session with a laser scanning confocal microscope (CarlZeiss LSM780, 40X ApoChromat, 1.4 NA, oil immersion objective). All images were collected as 0.2μm (for NMJ) or 0.1μm (for individual synapses) optical sections and the *z*-stacks were analyzed with Imaris software (Bitplane) or ImageJ (NIH) respectively.

NMJ morphometrics were performed as previously described (Ramos et al. 2015). Briefly, positive puncta were detected semi-automatically using the spot finding Imaris algorithm. To quantify fluorescence intensities, synaptic ROI areas surrounding anti-HRP immunoreactivities were selected and the signals measured individually at NMJs (muscle 6/7 or muscle 4, segment A3) from 10 or more different larvae for each genotype. The signal intensities were calculated relative to HRP volume and subsequently normalized to control. Morphometric quantifications such as branching points and branch length were quantified semi-automatically with Filament algorithm. Boutons were counted in preparations double labeled with anti-HRP and anti-Dlg; boutons volume were estimated by manual selection and Spot algorithm (Imaris). All quantifications were performed while blinded to genotype. Statistical analyses were performed using the Student t-test with a two-tailed distribution and a two-sample unequal variance. Error bars in all graphs indicate standard deviation ±SEM. ***; p<0.001, **; p<0.005, *; p<0.05, ns; p>0.05.

For the quantification of individual Brp puncta, singles NMJs (muscle 6/7, segment A3) stained for Brp and HRP were assembled from multiple frames (3-4) imaged at a 4.0x zoom. Individual frames were analyzed using ImageJ software for Fiji distribution and maximum intensity projections (Schindelin et al., 2012). The channels were separated and the low intensity Brp-positive removed by applying a threshold and a mask. The ‘RawIntDen’, which represent the total intensity of the Brp signal, and the ‘Area’ of the selection were calculated and added them together to assemble an entire NMJ from different frames. For each genotype, the Brp intensity per unit area (∑RawIntDen)/(∑Area) from PhTx treated animals was normalized and reported relative to the untreated larvae. Statistical analyses were performed with Prism7 using the unpaired t-test with a two-tailed distribution. For the Brp peak analysis, we measured the intensity of the peaks contained within the selected masks using the Find Maxima algoritm (ImageJ) and normalized them as above. The frequency distribution and cumulative distribution of peak values were calculated with Prism7.

### Super resolution (3D-SIM) imaging and data processing

Super-resolution imaging was performed on a Carl Zeiss Elyra PS1 inverted microscope using a Plan-Apo 100X (1.46 NA) oil immersion objective and an EM-CCD Andor iXon 885 camera. We collected ×5 phases at ×3 angles for a total of 15 images per plane. Singles NMJ 6/7 at the A3 segment were captured by multiple frames (3-4 per NMJ); the stacks of z-sections were taken at a spacing of every 100 nm. All raw images were processed and reconstructed in 3D using Zen Black 2010 software (Carl Zeiss). The images were also channel aligned using an alignment matrix generated by imaging colored beads. The PSD areas were estimated using the Fiji distribution algorithm (ImageJ) (Schindelin et al., 2012). Single ROIs corresponding to the maximum PSD areas were selected and measured using either the wand tool (with legacy and regulated tolerance) or manually, for overlapping regions. At least 1400 single PSDs from 12 or more different NMJs for each genotype. Statistical analyses were performed with Prism7 using One-way ANOVA Analysis with post-hoc Tukey test for multiple comparison, frequency distribution and cumulative distribution.

### Electrophysiology

The standard larval body wall muscle preparation first developed by Jan and Jan (1976) was used for electrophysiological recordings. Wandering third instar larvae were dissected and washed in physiological saline. Using a custom microscope stage system, all recordings were performed in HL-3 saline (Stewart et al., 1994) containing 0.5 mM CaCl_2_ unless otherwise indicated. For the acute homeostasis paradigm, semi-intact preparations were incubated with philanthotoxin-343 (PhTx) (Sigma; 20 μM) in Ca^2+^-free HL-3 saline for 15 min as previously described (Frank et al., 2006). The nerve roots were cut near the exiting site of the ventral nerve cord so that the motor nerve could be picked up by a suction electrode. Intracellular recordings were made from muscle 6, abdominal segment 3 and 4. Data were used when the input resistance of the muscle was >5 MΩ and the resting membrane potential was < −60 mV. The input resistance of the recording microelectrode (backfilled with 3 M KCl) ranged from 20 to 25 MΩ. Muscle synaptic potentials were recorded using Axon Clamp 2B amplifier (Axon Instruments) and analyzed using pClamp 10 software. Spontaneous miniature excitatory junction potentials (mEJPs) were recorded in the absence of any stimulation. To calculate mEJP mean amplitudes, 50–100 events from each muscle were measured and averaged using the Mini Analysis program (Synaptosoft). Minis with a slow rise and falling time arising from neighboring electrically coupled muscle cells were excluded from analysis. Evoked EJPs were recorded following supra-threshold stimuli (200 μsec) to the appropriate segmental nerve with a suction electrode. Ten to fifteen EJPs evoked by low frequency of stimulation (0.1 Hz) were averaged. Quantal content was calculated by dividing the mean EJP by the mean mEJP after correction of EJP amplitude for nonlinear summation according to previously described methods. Corrected EJP amplitude = E[Ln[E/(E - recorded EJP)]], where E is the difference between reversal potential and resting potential. The reversal potential used in this correction was 0 mV.

For readily released pool (RRP) measurements, evoked excitatory junction currents (EJCs) were recorded at a voltage clamped to – 65 mV, and 30 EJCs were stimulated at 50 Hz in HL-3 saline with 1.5 mM Ca^2+^ and 10 mM Mg^2+^. EJC amplitudes during a stimulus train were calculated by subtracting the baseline current just preceding an EJC from the subsequent peak of the EJC. The cumulative EJC amplitude was obtained by back-extrapolating a straight line fitted to the final 10 points of the cumulative EJC to time zero. The size of the RRP were calculated by dividing the cumulative EJC amplitude by the mean mEJP amplitude recorded in the same muscle. Statistical analysis used Prism7 using ANOVA followed by a Tukey post hoc test. Data are presented as mean ±SEM.

### Presynaptic Ca^2+^ imaging

Cytosolic Ca^2+^ levels were monitored through the fluorescence of a Ca^2+^-sensitive dye (Oregon-Green BAPTA-1; OGB-1) relative to a Ca^2+^-insensitive dye (Alexa Fluor 568; AF568); both of which were loaded into motor neuron terminals using the forward-filling technique as previously described (Macleod, 2012). Segment nerves were forward-filled with 10,000 MW dextran-conjugated OGB-1, in constant ratio with 10,000 MW dextran-conjugated AF568. Fluorescence imaging was performed through a water-dipping 100X 1.1 NA Nikon objective fitted to an upright Nikon Eclipse FN1 microscope. Fluorescence was excited using a Lumencor Spectra X light engine (OGB-1: 483/32 nm; AF568: 550/15 nm). Emitted light (OGB-1: 525/84 nm; AF568: 605/52 nm) was captured by an Andor iXon3 897 EMCCD camera running at 112 frames-per-second (2×2 binning, 8 ms exposures). OGB-1 images were not interdigitated with AF568 images during the stimulus protocol, rather, AF568 fluorescence images were captured immediately before and after the stimulus protocol to provide ratio information. While larvae were dissected and incubated in Schneider’s insect medium, this medium was replaced with HL3 at least 20 minutes prior to imaging. HL3 was supplemented with 0.5 mM Ca^2+^, 20 mM Mg^2+^, and 7 mM L-glutamic acid, which prevents muscle contraction (Macleod et al., 2004). Segmental nerves were stimulated according to the pattern illustrated in Figure 7A, where each fluorescence transient is the result of an impulse of approximately 1.5 volts applied to the nerve for 0.4 ms. The background fluorescence was subtracted from each image and the average pixel intensity was measured within a region-of-interest containing 2-5 non-terminal boutons using NIS-Elements AR software (Nikon). Fluorescence intensity traces were further processed in ImageJ [Fiji (fiji.sc; ImageJ)]. OGB-1 fluorescence was imaged for 5 seconds prior to the first stimulus pulse, and these data were used to estimate the OGB-1 bleach trend which was then numerically removed from the entire trace. Ca^2+^ levels are expressed as the fluorescence ratio of OGB-1 to AF568. Fluorescence transients corresponding to the action potentials evoked at 1 Hz were numerically averaged into a single trace and used to calculate peak amplitude and the decay time constant (τ). The amplitude of a single transient was calculated as the displacement between the baseline prior to the transient and the mono-exponential fit to the transient decay when extrapolated forward to the time of the nerve stimulus. Two criteria were used to exclude data from further analysis; first, when the data were collected from a terminal with a resting Ca^2+^ level that was assessed to be an outlier, and secondly, when single action potential evoked fluorescence transients did not recover to baseline with a time course of less than 150 ms. Outliers were defined using the median absolute deviation (MAD; (Leys et al., 2013)) were an outlier was considered to be any value beyond 3X MAD of the median. Differences between *neto-α ^null^* and control were tested using the Students T statistic, and where normality tests failed, the Mann Whitney U statistic was used.

### Electron microscopy

For transmission electron microscopy (TEM), *Drosophila* larva fillets were fixed in 2% glutaraldehyde/ 2% formaldehyde/2 mM CaCl2 in 0.1 M cacodylate buffer pH 7.4 for 15 min at room temperature followed by 1 hour on ice in fresh fixative. After 5 washes in the buffer, they were post-fixed in 2% osmium tetroxide in the same buffer for 2 hours on ice, washed once in the buffer and 5 times in double distilled water. The samples were than stained *en bloc* overnight in 2% aqueous uranyl acetate, washed 5x in water, dehydrated in series of ethanol concentrations and penetrated with EMbed 812 (EMS, Hatfield, PA). For easy orientation, the fillets were placed on a glass coverslip with the inside facing glass and embedded in the same resin subsequently polymerized at 65°C. The coverslip was removed using hydrofluoric acid; blocks containing fillets were cut out, re-mounted on holders inside facing out and cut parallel to the original glass surface. Semi-thin (200 nm) sections were cut, stained with toluidine blue and checked under light microscope. Once the exact position of cutting was reached, serial thin (80 nm) sections of the fillets were cut on Leica EM UC7 microtome (Leica, Deerfield, IL) and stained with uranyl acetate. The samples were examined on FEI Tecnai 20 TEM (FEI, Hillsboro OR) operated at 120 kV and images were recorded on AMT XR81 CCD camera (AMT, Woburn, MA). PSDs and T-bars metrics were quantified from 3-5 serial sections by selecting the maximum PSD length, T-bar platform and pedestal.

## Acknowledgments

T.H.H, R.V., C.I.R., Q.W., P.N., and M.S. were supported by Intramural Program of the National Institutes of Health, *Eunice Kennedy Shriver* National Institute of Child Health and Human Development, grants ZIA HD008914 and ZIA HD008869 awarded to M.S. G.T.M., R.X.H. and M.S. were supported by NIH NINDS award NS061914. We thank Chi-Hon Lee and Tom Brody for comments and discussions on this manuscript. We also thank the Bloomington Stock Center at Indiana University for fly stocks and the Developmental Studies Hybridoma Bank at the University of Iowa for antibodies.

## Author Contributions

Conceptualization, T.H.H., C.I.R., R.V., G.M. and M.S.; Methodology, T.H.H., C.I.R., R.V., G.M. and M.S.; Investigation, T.H.H., C.I.R., R.V., Q.W., M.J., M.S., R.H., G.M. and M.S.; Writing, T.H.H., C.I.R., R.V., G.M. and M.S.; Funding Acquisition, G.M. and M.S., Resources, P.N. and M.L.; Supervision, G.M. and M.S.

---

**Supplemental Table 1:**
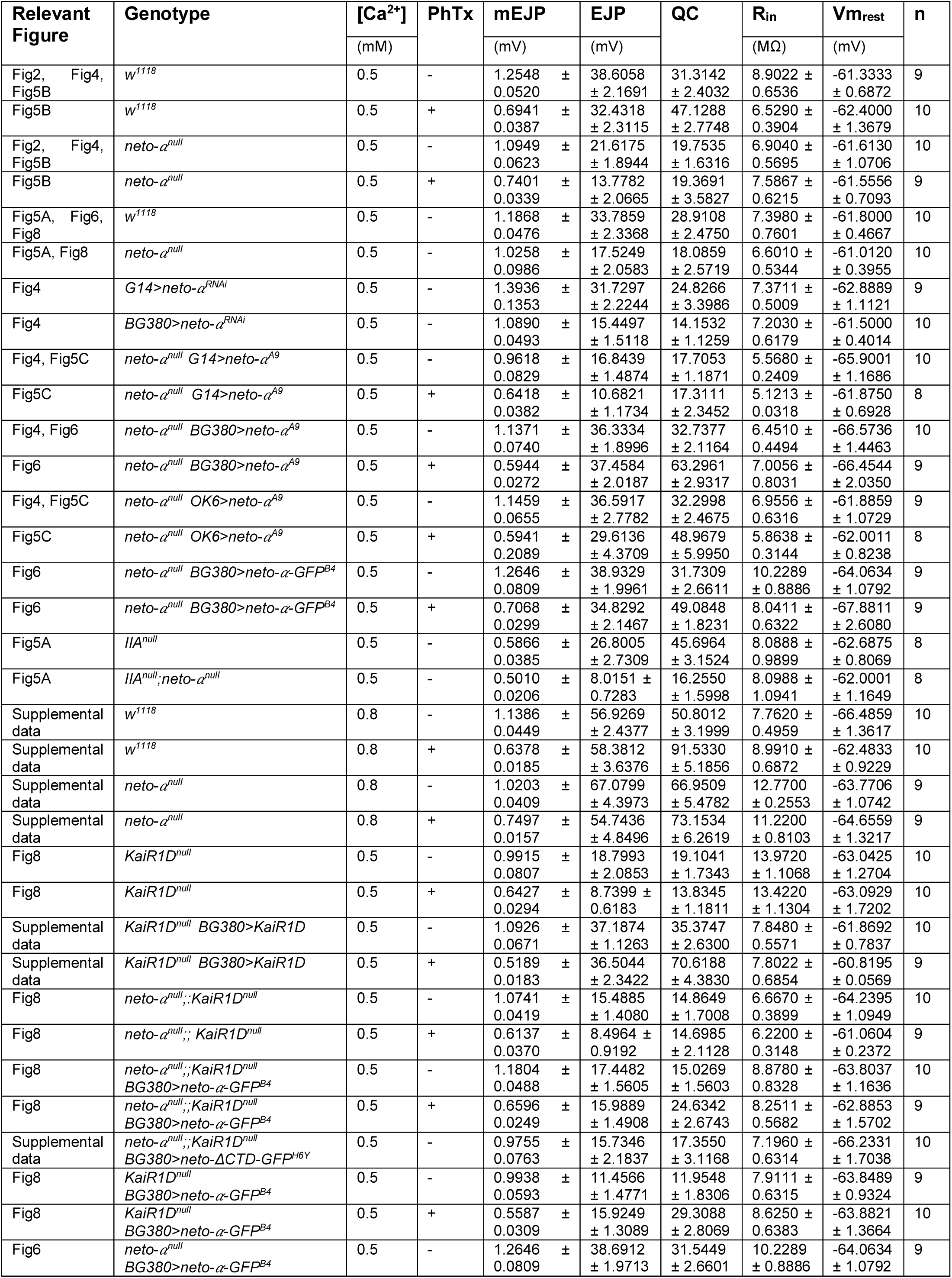
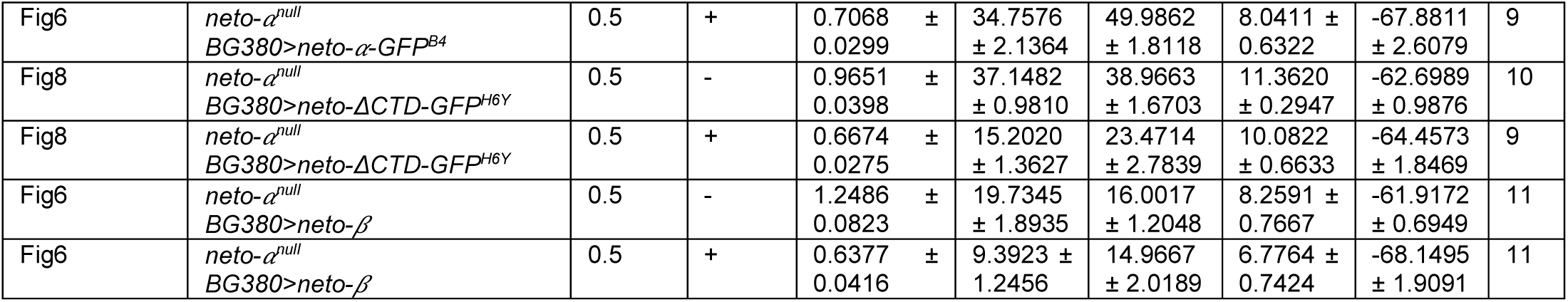

## References

Alberstein, R., Grey, R., Zimmet, A., Simmons, D.K., and Mayer, M.L. (2015). Glycine activated ion channel subunits encoded by ctenophore glutamate receptor genes. Proc Natl Acad Sci U S A 112, E6048–6057.

Banovic, D., Khorramshahi, O., Owald, D., Wichmann, C., Riedt, T., Fouquet, W., Tian, R., Sigrist, S.J., and Aberle, H. (2010). Drosophila neuroligin 1 promotes growth and postsynaptic differentiation at glutamatergic neuromuscular junctions. Neuron 66, 724–738.

Copits, B.A., and Swanson, G.T. (2012). Dancing partners at the synapse: auxiliary subunits that shape kainate receptor function. Nat Rev Neurosci 13, 675–686.

Davis, G.W., DiAntonio, A., Petersen, S.A., and Goodman, C.S. (1998). Postsynaptic PKA controls quantal size and reveals a retrograde signal that regulates presynaptic transmitter release in Drosophila. Neuron 20, 305–315.

Davis, G.W., and Muller, M. (2015). Homeostatic control of presynaptic neurotransmitter release. Annual review of physiology 77, 251–270.

DiAntonio, A. (2006). Glutamate receptors at the Drosophila neuromuscular junction. Int Rev Neurobiol 75, 165–179.

DiAntonio, A., Petersen, S.A., Heckmann, M., and Goodman, C.S. (1999). Glutamate receptor expression regulates quantal size and quantal content at the Drosophila neuromuscular junction. J Neurosci 19, 3023–3032.

Dickman, D.K., and Davis, G.W. (2009). The schizophrenia susceptibility gene dysbindin controls synaptic homeostasis. Science 326, 1127–1130.

Dickman, D.K., Tong, A., and Davis, G.W. (2012). Snapin is critical for presynaptic homeostatic plasticity. J Neurosci 32, 8716–8724.

Featherstone, D.E., Rushton, E., Rohrbough, J., Liebl, F., Karr, J., Sheng, Q., Rodesch, C.K., and Broadie, K. (2005). An essential Drosophila glutamate receptor subunit that functions in both central neuropil and neuromuscular junction. J Neurosci 25, 3199–3208.

Fouquet, W., Owald, D., Wichmann, C., Mertel, S., Depner, H., Dyba, M., Hallermann, S., Kittel, R.J., Eimer, S., and Sigrist, S.J. (2009). Maturation of active zone assembly by Drosophila Bruchpilot. J Cell Biol 186, 129–145.

Frank, C.A. (2014). Homeostatic plasticity at the Drosophila neuromuscular junction. Neuropharmacology 78, 63–74.

Frank, C.A., Kennedy, M.J., Goold, C.P., Marek, K.W., and Davis, G.W. (2006). Mechanisms underlying the rapid induction and sustained expression of synaptic homeostasis. Neuron 52, 663–677.

Frank, C.A., Pielage, J., and Davis, G.W. (2009). A presynaptic homeostatic signaling system composed of the Eph receptor, ephexin, Cdc42, and CaV2.1 calcium channels. Neuron 61, 556–569.

Goel, P., Li, X., and Dickman, D. (2017). Disparate Postsynaptic Induction Mechanisms Ultimately Converge to Drive the Retrograde Enhancement of Presynaptic Efficacy. Cell reports 21, 2339–2347.

Goold, C.P., and Davis, G.W. (2007). The BMP ligand Gbb gates the expression of synaptic homeostasis independent of synaptic growth control. Neuron 56, 109–123.

Gratz, S.J., Rubinstein, C.D., Harrison, M.M., Wildonger, J., and O’Connor-Giles, K.M. (2015). CRISPR-Cas9 Genome Editing in Drosophila. Curr Protoc Mol Biol 111, 31 32 31–20.

Guan, B., Hartmann, B., Kho, Y.H., Gorczyca, M., and Budnik, V. (1996). The Drosophila tumor suppressor gene, dlg, is involved in structural plasticity at a glutamatergic synapse. Curr Biol 6, 695–706.

Han, T.H., Dharkar, P., Mayer, M.L., and Serpe, M. (2015). Functional reconstitution of Drosophila melanogaster NMJ glutamate receptors. Proc Natl Acad Sci U S A 112, 6182–6187.

Jackson, A.C., and Nicoll, R.A. (2011). The expanding social network of ionotropic glutamate receptors: TARPs and other transmembrane auxiliary subunits. Neuron 70, 178–199.

Jan, L.Y., and Jan, Y.N. (1982). Antibodies to horseradish peroxidase as specific neuronal markers in Drosophila and in grasshopper embryos. Proc Natl Acad Sci U S A 79, 2700–2704.

Kim, Y.J., Bao, H., Bonanno, L., Zhang, B., and Serpe, M. (2012). Drosophila Neto is essential for clustering glutamate receptors at the neuromuscular junction. Genes Dev 26, 974–987.

Kim, Y.J., Igiesuorobo, O., Ramos, C.I., Bao, H., Zhang, B., and Serpe, M. (2015). Prodomain removal enables neto to stabilize glutamate receptors at the Drosophila neuromuscular junction. PLoS genetics 11, e1004988.

Kim, Y.J., and Serpe, M. (2013). Building a synapse: A complex matter. Fly 7.

Kiragasi, B., Wondolowski, J., Li, Y., and Dickman, D.K. (2017). A Presynaptic Glutamate Receptor Subunit Confers Robustness to Neurotransmission and Homeostatic Potentiation. Cell reports 19, 2694–2706.

Kittel, R.J., Wichmann, C., Rasse, T.M., Fouquet, W., Schmidt, M., Schmid, A., Wagh, D.A., Pawlu, C., Kellner, R.R., Willig, K.I., et al. (2006). Bruchpilot promotes active zone assembly, Ca2+ channel clustering, and vesicle release. Science 312, 1051–1054.

Lerma, J., and Marques, J.M. (2013). Kainate receptors in health and disease. Neuron 80, 292–311.

Letts, V.A., Felix, R., Biddlecome, G.H., Arikkath, J., Mahaffey, C.L., Valenzuela, A., Bartlett, F.S., 2nd, Mori, Y., Campbell, K.P., and Frankel, W.N. (1998). The mouse stargazer gene encodes a neuronal Ca2+-channel gamma subunit. Nat Genet 19, 340–347.

Leys, C., Ley, C., Klein, O., Bernard, P., and Licata, L. (2013). Detecting outliers: Do not use standard deviation around the mean, use absolute deviation around the median. Journal of Experimental Social Psychology 49, 764–766.

Li, X., Goel, P., Chen, C., Angajala, V., Chen, X., and Dickman, D.K. (2018). Synapse-specific and compartmentalized expression of presynaptic homeostatic potentiation. Elife 7.

Li, Y., Dharkar, P., Han, T.H., Serpe, M., Lee, C.H., and Mayer, M.L. (2016). Novel Functional Properties of Drosophila CNS Glutamate Receptors. Neuron 92, 1036–1048.

Littleton, J.T., and Ganetzky, B. (2000). Ion channels and synaptic organization: analysis of the Drosophila genome. Neuron 26, 35–43.

Lomash, R.M., Sheng, N., Li, Y., Nicoll, R.A., and Roche, K.W. (2017). Phosphorylation of the kainate receptor (KAR) auxiliary subunit Neto2 at serine 409 regulates synaptic targeting of the KAR subunit GluK1. J Biol Chem 292, 15369–15377.

Macleod, G.T. (2012). Forward-filling of dextran-conjugated indicators for calcium imaging at the Drosophila larval neuromuscular junction. Cold Spring Harb Protoc 2012, 791–796.

Macleod, G.T., Marin, L., Charlton, M.P., and Atwood, H.L. (2004). Synaptic vesicles: test for a role in presynaptic calcium regulation. J Neurosci 24, 2496–2505.

Marrus, S.B., Portman, S.L., Allen, M.J., Moffat, K.G., and DiAntonio, A. (2004). Differential localization of glutamate receptor subunits at the Drosophila neuromuscular junction. J Neurosci 24, 1406–1415.

Mayer, M.L. (2017). The Challenge of Interpreting Glutamate-Receptor Ion-Channel Structures. Biophysical journal 113, 2143–2151.

Milstein, A.D., and Nicoll, R.A. (2008). Regulation of AMPA receptor gating and pharmacology by TARP auxiliary subunits. Trends Pharmacol Sci 29, 333–339.

Mosca, T.J., Hong, W., Dani, V.S., Favaloro, V., and Luo, L. (2012). Trans-synaptic Teneurin signalling in neuromuscular synapse organization and target choice. Nature 484, 237–241.

Muller, M., and Davis, G.W. (2012). Transsynaptic control of presynaptic Ca(2)(+) influx achieves homeostatic potentiation of neurotransmitter release. Curr Biol 22, 1102–1108.

Muller, M., Liu, K.S., Sigrist, S.J., and Davis, G.W. (2012). RIM controls homeostatic plasticity through modulation of the readily-releasable vesicle pool. J Neurosci 32, 16574–16585.

Muller, M., Pym, E.C., Tong, A., and Davis, G.W. (2011). Rab3-GAP controls the progression of synaptic homeostasis at a late stage of vesicle release. Neuron 69, 749–762.

Ng, D., Pitcher, G.M., Szilard, R.K., Sertie, A., Kanisek, M., Clapcote, S.J., Lipina, T., Kalia, L.V., Joo, D., McKerlie, C., et al. (2009). Neto1 is a novel CUB-domain NMDA receptor-interacting protein required for synaptic plasticity and learning. PLoS Biol 7, e41.

Orr, B.O., Fetter, R.D., and Davis, G.W. (2017). Retrograde semaphorin-plexin signalling drives homeostatic synaptic plasticity. Nature 550, 109–113.

Perrais, D., Veran, J., and Mulle, C. (2010). Gating and permeation of kainate receptors: differences unveiled. Trends Pharmacol Sci 31, 516–522.

Petersen, S.A., Fetter, R.D., Noordermeer, J.N., Goodman, C.S., and DiAntonio, A. (1997). Genetic analysis of glutamate receptors in Drosophila reveals a retrograde signal regulating presynaptic transmitter release. Neuron 19, 1237–1248.

Petzoldt, A.G., Lee, Y.H., Khorramshahi, O., Reynolds, E., Plested, A.J., Herzel, H., and Sigrist, S.J. (2014). Gating characteristics control glutamate receptor distribution and trafficking in vivo. Curr Biol 24, 2059–2065.

Pinheiro, P.S., Perrais, D., Coussen, F., Barhanin, J., Bettler, B., Mann, J.R., Malva, J.O., Heinemann, S.F., and Mulle, C. (2007). GluR7 is an essential subunit of presynaptic kainate autoreceptors at hippocampal mossy fiber synapses. Proc Natl Acad Sci U S A 104, 12181–12186.

Qin, G., Schwarz, T., Kittel, R.J., Schmid, A., Rasse, T.M., Kappei, D., Ponimaskin, E., Heckmann, M., and Sigrist, S.J. (2005). Four different subunits are essential for expressing the synaptic glutamate receptor at neuromuscular junctions of Drosophila. J Neurosci 25, 3209–3218.

Ramos, C.I., Igiesuorobo, O., Wang, Q., and Serpe, M. (2015). Neto-mediated intracellular interactions shape postsynaptic composition at the Drosophila neuromuscular junction. PLoS genetics 11, e1005191.

Ren, X., Sun, J., Housden, B.E., Hu, Y., Roesel, C., Lin, S., Liu, L.P., Yang, Z., Mao, D., Sun, L., et al. (2013). Optimized gene editing technology for Drosophila melanogaster using germ line-specific Cas9. Proc Natl Acad Sci U S A 110, 19012–19017.

Schindelin, J., Arganda-Carreras, I., Frise, E., Kaynig, V., Longair, M., Pietzsch, T., Preibisch, S., Rueden, C., Saalfeld, S., Schmid, B., et al. (2012). Fiji: an open-source platform for biological-image analysis. Nature methods 9, 676–682.

Sheng, N., Bemben, M.A., Diaz-Alonso, J., Tao, W., Shi, Y.S., and Nicoll, R.A. (2018). LTP requires postsynaptic PDZ-domain interactions with glutamate receptor/auxiliary protein complexes. Proc Natl Acad Sci U S A 115, 3948–3953.

Stewart, B.A., Atwood, H.L., Renger, J.J., Wang, J., and Wu, C.F. (1994). Improved stability of Drosophila larval neuromuscular preparations in haemolymph-like physiological solutions. J Comp Physiol A 175, 179–191.

Straub, C., Hunt, D.L., Yamasaki, M., Kim, K.S., Watanabe, M., Castillo, P.E., and Tomita, S. (2011). Distinct functions of kainate receptors in the brain are determined by the auxiliary subunit Neto1. Nat Neurosci 14, 866–873.

Sulkowski, M.J., Han, T.H., Ott, C., Wang, Q., Verheyen, E.M., Lippincott-Schwartz, J., and Serpe, M. (2016). A Novel, Noncanonical BMP Pathway Modulates Synapse Maturation at the Drosophila Neuromuscular Junction. PLoS genetics 12, e1005810.

Sumioka, A., Yan, D., and Tomita, S. (2010). TARP phosphorylation regulates synaptic AMPA receptors through lipid bilayers. Neuron 66, 755–767.

Tang, M., Ivakine, E., Mahadevan, V., Salter, M.W., and McInnes, R.R. (2012). Neto2 Interacts with the Scaffolding Protein GRIP and Regulates Synaptic Abundance of Kainate Receptors. PLoS One 7, e51433.

Tomita, S. (2010). Regulation of ionotropic glutamate receptors by their auxiliary subunits. Physiology (Bethesda) 25, 41–49.

Tomita, S., Adesnik, H., Sekiguchi, M., Zhang, W., Wada, K., Howe, J.R., Nicoll, R.A., and Bredt, D.S. (2005). Stargazin modulates AMPA receptor gating and trafficking by distinct domains. Nature 435, 1052–1058.

Tomita, S., and Castillo, P.E. (2012). Neto1 and Neto2: auxiliary subunits that determine key properties of native kainate receptors. J Physiol 590, 2217–2223.

Tomita, S., Chen, L., Kawasaki, Y., Petralia, R.S., Wenthold, R.J., Nicoll, R.A., and Bredt, D.S. (2003). Functional studies and distribution define a family of transmembrane AMPA receptor regulatory proteins. J Cell Biol 161, 805–816.

Twomey, E.C., Yelshanskaya, M.V., Grassucci, R.A., Frank, J., and Sobolevsky, A.I. (2016). Elucidation of AMPA receptor-stargazin complexes by cryo-electron microscopy. Science 353, 83–86.

Wagh, D.A., Rasse, T.M., Asan, E., Hofbauer, A., Schwenkert, I., Durrbeck, H., Buchner, S., Dabauvalle, M.C., Schmidt, M., Qin, G., et al. (2006). Bruchpilot, a protein with homology to ELKS/CAST, is required for structural integrity and function of synaptic active zones in Drosophila. Neuron 49, 833–844.

Walker, C.S., Brockie, P.J., Madsen, D.M., Francis, M.M., Zheng, Y., Koduri, S., Mellem, J.E., Strutz-Seebohm, N., and Maricq, A.V. (2006). Reconstitution of invertebrate glutamate receptor function depends on stargazin-like proteins. Proc Natl Acad Sci U S A 103, 10781–10786.

Wang, Q., Chiu, S.L., Koropouli, E., Hong, I., Mitchell, S., Easwaran, T.P., Hamilton, N.R., Gustina, A.S., Zhu, Q., Ginty, D.D., et al. (2017). Neuropilin-2/PlexinA3 Receptors Associate with GluA1 and Mediate Sema3F-Dependent Homeostatic Scaling in Cortical Neurons. Neuron 96, 1084–1098 e1087.

Wang, R., Mellem, J.E., Jensen, M., Brockie, P.J., Walker, C.S., Hoerndli, F.J., Hauth, L., Madsen, D.M., and Maricq, A.V. (2012). The SOL-2/Neto auxiliary protein modulates the function of AMPA-subtype ionotropic glutamate receptors. Neuron 75, 838–850.

Wang, T., Jones, R.T., Whippen, J.M., and Davis, G.W. (2016). alpha2delta-3 Is Required for Rapid Transsynaptic Homeostatic Signaling. Cell reports 16, 2875–2888.

Weyhersmuller, A., Hallermann, S., Wagner, N., and Eilers, J. (2011). Rapid active zone remodeling during synaptic plasticity. J Neurosci 31, 6041–6052.

Wyeth, M.S., Pelkey, K.A., Yuan, X., Vargish, G., Johnston, A.D., Hunt, S., Fang, C., Abebe, D., Mahadevan, V., Fisahn, A., et al. (2017). Neto Auxiliary Subunits Regulate Interneuron Somatodendritic and Presynaptic Kainate Receptors to Control Network Inhibition. Cell reports 20, 2156–2168.

Xing, G., Gan, G., Chen, D., Sun, M., Yi, J., Lv, H., Han, J., and Xie, W. (2014). Drosophila neuroligin3 regulates neuromuscular junction development and synaptic differentiation. J Biol Chem 289, 31867–31877.

Zhang, W., St-Gelais, F., Grabner, C.P., Trinidad, J.C., Sumioka, A., Morimoto-Tomita, M., Kim, K.S., Straub, C., Burlingame, A.L., Howe, J.R., et al. (2009). A transmembrane accessory subunit that modulates kainate-type glutamate receptors. Neuron 61, 385–396.

